# EMC holdase:Ca_V_1.2/Ca_V_β_3_ complex and Ca_V_1.2 channel structures reveal Ca_V_ assembly and drug binding mechanisms

**DOI:** 10.1101/2022.10.03.510667

**Authors:** Zhou Chen, Abhisek Mondal, Fayal Abderemane-Ali, José Montano, Balyn Zaro, Daniel L. Minor

## Abstract

Voltage-gated ion channels (VGICs) comprise multiple structural units whose assembly is required for function^1,2^. There is scant structural understanding of how VGIC subunits assemble and whether chaperone proteins are required. High-voltage activated calcium channels (Ca_V_s)^3,4^ are paradigmatic multi-subunit VGICs from electrically excitable tissues whose function and trafficking is powerfully shaped by interactions between pore-forming Ca_V_1 or Ca_V_2 Ca_V_α_1^3^_ and auxiliary Ca_V_β_5_, and Ca_V_α_2_δ subunits^6,7^. Here, we present cryo-EM structures of human brain and cardiac Ca_V_1.2 bound with Ca_V_β_3_ to a chaperone, the endoplasmic reticulum membrane protein complex (EMC)^8,9^, and of the isolated Ca_V_1.2/Ca_V_β_3_/Ca_V_α_2_δ-1 channel. These provide an unprecedented view of an EMC holdase:client complex and define EMC sites, the TM and Cyto docks, whose interaction with the client channel cause partial extraction of a pore subunit and splay open the Ca_V_α_2_δ interaction site. The structures further identify the Ca_V_α_2_δ binding site for gabapentinoid anti-pain and anti-anxiety drugs^6^, show that EMC and Ca_V_α_2_δ channel interactions are mutually exclusive, and indicate that EMC to Ca_V_α_2_δ handoff involves a Ca^2+^-dependent step and ordering of multiple Ca_V_1.2 elements. Together, the structures unveil a Ca_V_ assembly intermediate and previously unknown EMC client binding sites that have broad implications for biogenesis of VGICs and other membrane proteins.

## Introduction

All voltage gated ion channel (VGIC) superfamily members require assembly to make the functional channel^1,2^. Voltage-gated calcium channels (Ca_V_s) from the high voltage activated subfamilies (Ca_V_1 and Ca_V_2)^3,4^ are ubiquitous in electrically excitable tissues such as muscle, the brain, and heart and provide a paradigmatic example of a multi-subunit VGIC comprising three key components, the pore-forming Ca_V_α_1_, cytoplasmic Ca_V_β^5^, and extracellular Ca_V_α_2_δ ^6,7^ subunits. It has been known for more than 30 years that Ca_V_β and Ca_V_α_2_δ association with Ca_V_α_1_ shape Ca_V_ biophysical properties and plasma membrane expression in profound ways^5,10–13^. Ca_V_α_2_δ has a particularly important role in enhancing cell surface expression and is the target of widely used anti-nociceptive and anti-anxiety drugs from the gabapentinoid class^13–15^ that bind Ca_V_α_2_δ and impair surface expression^14,16,17^.

The importance of subunit assembly for biogenesis of properly functioning channels has been long appreciated^2,18^. Yet, there remains scarce understanding of the steps involved in this process or the extent to which chaperone proteins may participate. The EMC (ER membrane protein complex) is a nine protein transmembrane assembly thought to aid the membrane insertion of tail anchored proteins^8,9,19,20^ and transmembrane segments having mixed hydrophobic/hydrophilic character such as those in ion channels^21–23^, receptors^24–26^, and transporters^22,24,27^. It may also function as holdase for partially assembled membrane proteins^26^. Although recent structural studies have outlined the EMC architecture and have implicated various EMC client protein interaction sites based on mutational analysis^26,28–30^, there are no structural examples of EMC:client protein complexes that would guide understanding of how the EMC engages substrate proteins.

Here, we present cryo-EM structures of a ~0.6 MDa complex containing the human brain and heart Ca_V_1.2 Ca_V_α_1_ subunit^3^, CaVβ_3_, and human EMC and the fully-assembled Ca_V_1.2/Ca_V_β_3_/Ca_v_α_2_δ-1 channel having overall resolutions of 3.4Å and 3.3Å, respectively. This Ca_V_ is the target of numerous drugs for cardiac and neurological diseases^6,7,31^ and the object of many arrhythmia, autism, and neuropsychiatric disease mutations^3,4,32^. Our data provide an unprecedented view of an EMC:client membrane protein and reveal EMC sites, the TM and Cyto docks, whose interaction with the Ca_V_α:Ca_V_β complex results in a partial extraction of one of the channel pore domains (PDs) and a splaying open of the extracellular face where Ca_V_α_2_δ binds in the fully assembled channel. Together, the structural data indicate that EMC and Ca_V_α_2_δ binding to the Ca_V_α:Ca_V_β channel core is mutually exclusive and that EMC handoff to Ca_V_α_2_δ involves ordering three extracellular sites that are splayed apart in the EMC holdase complex. The Ca_V_1.2/Ca_V_β_3_/Ca_v_α_2_δ-1 structure also reveals a bound leucine that identifies the Ca_V_α_2_δ binding site targeted by widely-used gabapentinoid drugs for pain, fibromyalgia, postherpetic neuralgia, and generalized anxiety disorder^6,7^. Our data define a Ca_V_ assembly intermediate that establishes a new framework for understanding Ca_V_ biogenesis, drug action^3,6,33^, and consequences of Ca_V_ disease mutations affecting the brain, heart, and endocrine systems^4,32^. The newly discovered EMC:client interaction sites are likely to be exploited by many VGIC superfamily client proteins as well as other membrane protein classes.

## Results

### Discovery of a stable ER chaperone:ion channel complex

Purification of Ca_V_1.2(ΔC), a recombinantly-expressed human Ca_V_1.2 construct bearing a C-terminal truncation distal to the IQ domain having functional properties equivalent to full length Ca_V_1.2 (Fig. S1) (186 kDa), with rabbit Ca_V_β_3_ (54 kDa), yielded a sample displaying a good size exclusion profile that contained numerous unexpected bands (Figs. S2a-b). Mass spectrometry analysis showed that these bands comprise the nine subunits of a ~300 kDa membrane endoplasmic reticulum (ER) chaperone, the ER membrane protein complex (EMC)^8,9,19^, that co-purified with Ca_V_1.2(ΔC)/Ca_V_β_3_ (Table S1). Although EMC function is poorly understood^9,34^, this assembly is thought to aid membrane insertion of tail anchored proteins^20^ and transmembrane segments having mixed hydrophobic/hydrophilic character such as those found in ion channels^21–23^, receptors^24–26^, and transporters^22,24,27^, and may also function as holdase for partially assembled membrane proteins^26^. Cryo-EM images showed good quality particles that yielded two-dimensional class averages of a transmembrane complex having an unmistakable extramembrane EMC lumenal domain crescent (Figs. S2c-d). Co-expression of Ca_V_1.2(ΔC)/Ca_V_β_3_ with rabbit Ca_v_β_2_δ-1 (125 kDa) yielded a similarly behaved sample containing Ca_V_1.2(ΔC), Ca_V_β_3_, Ca_V_α_2_δ-1, and all nine EMC components (Figs. S2e-f) that also produced good cryo-EM images (Figs. S2g-h). Single particle analysis of both datasets (Fig. S2i) identified EMC:Ca_V_1.2(ΔC)/Ca_V_β_3_ complexes in both and Ca_V_1.2(ΔC)/Ca_V_β_3_/Ca_V_α_2_δ-1 complexes in the Ca_V_1.2(ΔC)/Ca_V_β_3_/Ca_V_α_2_δ-1 sample. The resultant EMC:Ca_V_1.2(ΔC)/Ca_V_β_3_ and Ca_V_1.2(ΔC)/Ca_V_β_3_/Ca_V_α_2_δ-1 maps had overall resolutions of 3.4Å and 3.3Å (Figs. S3a-b) and local resolutions as good as 2.4Å and 2.0Å, respectively, (Figs. S3c-f and S4-S7, Table S2). The maps clearly defined the subunits of the eleven protein EMC:Ca_V_1.2(ΔC)/Ca_V_β_3_ complex (Figs. S4a-b) and the tripartite Ca_V_1.2(ΔC)/Ca_V_β_3_/Ca_V_α_2_δ-1 channel assembly (Figs. S4c-d).

### Structure of the EMC:Ca_V_1.2/Ca_V_β_3_ complex

The EMC:Ca_V_1.2/Ca_V_β_3_ complex comprises a ~0.6 MDa assembly having dimensions of ~220Å normal to and ~100 × 130Å parallel to the membrane plane, respectively (Fig. 1a). Structural analysis identifies extensive Ca_V_1.2 and Ca_V_β_3_ interactions with the EMC (Fig. 1a) comprising two binding sites, the ‘TM dock’ and ‘Cyto dock’. These two sites are ~90° apart from each other with respect to the EMC core and engage transmembrane and cytoplasmic elements of the Ca_V_1.2/Ca_V_β_3_ channel complex, respectively (Figs. 1b-c, Movie S1).

**Figure 1.**
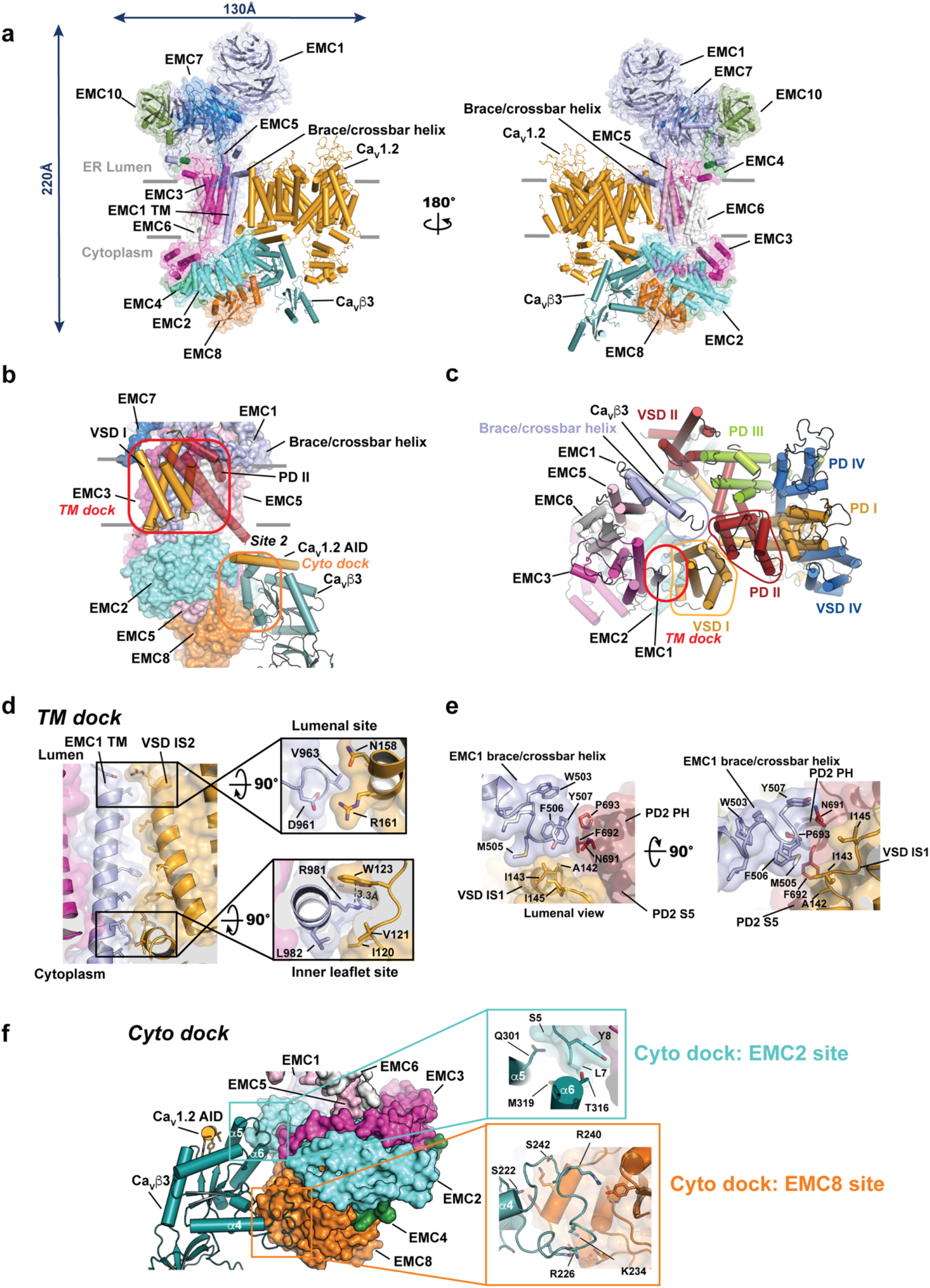
Structure of the EMC:Ca_V_1.2(ΔC)/Ca_V_β_3_ complex. **a**, Cartoon structure side views of the EMC:Ca_V_1.2(ΔC)/Ca_V_β_3_ complex. EMC1 (light blue), EMC2 (aquamarine), EMC3 (light magenta), EMC4 (forest), EMC5 (light pink), EMC6 (white), EMC7 (marine), EMC8 (orange), EMC10 (smudge), Ca_V_1.2 (bright orange), and Ca_V_β_3_ (light teal). EMC subunits are shown with semi-transparent surfaces. Dark blue arrows indicate dimensions. **b**, Location of TM dock and Cyto dock EMC client binding sites. Ca_V_1.2 components interacting with the EMC are shown as cylinders and colored: VSD I (yellow orange), PD II (firebrick), and Ca_V_β_3_ (light teal). AID helix (yellow orange) is also shown. EMC subunit colors are as in ‘**a**’. Grey bars in ‘a’ and ‘b’ denote the membrane. **c**, Lumenal view of the EMC:Ca_V_1.2(ΔC)/Ca_V_β_3_ complex transmembrane elements. EMC components are shown as cylinders and colored as in ‘**a**’. Ca_V_1.2(ΔC) domains are colored as Domain I (yellow orange), Domain II (firebrick), Domain III (lime), and Domain IV (marine). VSD and PD elements are labeled. Red oval shows TM dock site. Light blue circle shows the site of interaction of the EMC1 brace/crossbar helix with Ca_V_1.2 VSD I and PD II (outlined). **d**, Side view of the TM dock:VSD I interaction. Insets show details of interactions in the Lumenal site and Inner leaflet side. EMC1 (light blue), Ca_V_1.2 VSD I (yellow orange). **e**, Details of the EMC1 brace/crossbar helix with Ca_V_1.2 VSD I (yellow orange) and PD II (firebrick). **f**, Details of the Cyto dock. Ca_V_β_3_ (light teal) and AID helix (yellow orange) are shown as cylinders. EMC2 (aquamarine), EMC3 (light magenta), EMC4 (forest), EMC5 (light pink), EMC6 (white), and EMC8 (orange) are shown as surfaces. Insets show details of the EMC2 and EMC8 sites.

The TM dock comprises a large interface (1051 Å^2^) in which the single EMC1 transmembrane helix (EMC1 TM) binds the groove between the IS1 and IS2 transmembrane helices of the first Ca_V_1.2 voltage-sensor domain (VSD I) (Figs. 1b-d, S8a-b). There are extensive van der Waals interactions along the entire interface (Figs. S8a-b) that are framed by lumenal and inner bilayer leaflet hydrophilic interaction sites (Fig. 1d). The lumenal site involves a salt bridge between EMC1 Asp961, a conserved residue implicated in interacting with client proteins having hydrophilic character^30^, and Ca_V_1.2 VSD I Arg161, a residue found in all Ca_V_1s except Ca_V_1.4, as well as in Ca_V_2.3 (Fig. S8c). The inner leaflet site entails a cation-π interaction^35^ between the EMC1 Arg981 guanidinium sidechain and the Ca_V_1.2 IS1 Trp123 indole that with IS1 Ile120 frames a hydrophobic pocket, the ‘cation-π pocket’ (Figs. S8a-b). Preservation of these two hydrophilic interaction sites and the IS1/IS2 surface among Ca_V_1s and Ca_V_2s (Fig. S8c) suggests that other Ca_V_s VSD Is interact with the EMC similarly. In addition to these EMC1 TM interactions, the interfacial EMC1 helix (termed the ‘brace’^26^ or ‘crossbar’ helix^28^) extends its C-terminal end to contact the interface between VSD I and pore domain II (PD II) of the channel (Figs. 1c and e).

The Cyto dock is composed entirely of interactions between the EMC cytoplasmic cap, comprising EMC2, EMC3, EMC4, EMC5, and EMC8, and Ca_V_β_3_ (Fig. 1f), and involves two sites. In the larger site (the ‘EMC8 site’, 962 Å^2^), two Ca_V_β nucleotide kinase (NK) domain loops (the β7-β4 loop Thr218-Ala243 and β8-β9 loop Pro277-Lys282) bind EMC8 at a site whose center is formed by the last EMC8 helix (Fig. 1f). This interaction involves network of hydrogen bonds and salt bridges formed by seven conserved Ca_V_β residues (Ca_V_β:*EMC*8 pairs: Asp220:*His208*, Ser222:*His208*, Arg226:*Glu166*, Lys234:*Thr8*, Arg240:*Asp56/Tyr87*, Ser242:*Lys204*, and Gln279:*His208*) (Figs. 1f, and S8d-f) and includes a region, Lys225-Ser245, that is disordered in the isolated Ca_V_β_3_^36^ (Fig. S8g) but that becomes ordered upon EMC8 binding, suggesting that its conservation is linked to its role in EMC binding. In the smaller site (the ‘EMC2 site’, 550 Å^2^), Ca_V_β Gln301, Thr316, and Met319 contact the EMC2 N-terminal region through van der Waals interactions (Figs. 1f and S8 f-h). This site involves the Ca_V_β α5 and α6 helices that are part of the α-binding pocket (ABP) where all Ca_V_βs make high affinity interactions to the Ca_V_β_1_ subunit β-interaction domain (AID)^37,38^. The shared architecture of Ca_V_β subunits^37,38^ and strong conservation Ca_V_β contact residues indicate that all Ca_V_β isoforms should bind the Cyto dock.

Although the TM-dock site agrees with functional data implicating EMC1 Asp961 in client protein interaction^30^, the EMC1 TM:Ca_V_1.2 VSD I interface is on the opposite side of the EMC3, EMC5, and EMC6 gated transmembrane cavity proposed to act in membrane protein insertion^26,29,39^. The side of the channel facing the brace/crossbar helix surrounds the inner, hydrophobic, lipid-filled cavity formed by EMC1, EMC3, EMC5, and EMC6^26,28,29,39^, but apart from the interactions with the EMC1 brace/crossbar helix, there are no interactions with elements of this cavity. The Cyto dock site is on the opposite side of the membrane from the lumenal domain elements in EMC1, EMC7, and EMC10 suggested have a holdase function^26^. Further, the EMC2 and EMC8 sites are not near any of the cytoplasmic cap elements proposed for client interactions^28^. Hence, our structure presents a novel mechanism by which the EMC engages client proteins. Importantly, Ca_V_1.2/Ca_V_β_3_ interactions with the TM dock and Cyto dock involve highly conserved channel subunit elements (Figs. S8c and f), suggesting that Ca_V_s comprising various Ca_V_α_1_/Ca_V_α combinations^3,5^ can make the observed EMC:client interactions.

### Ca_V_1.2/Ca_V_β_3_/Ca_v_α_2_δ-1 structure identifies a Ca_V_α_2_δ drug binding site and a blocking lipid

The Ca_V_1.2(ΔC)/Ca_V_β_3_/Ca_v_α_2_δ-1 structure (Figs. 2a, S4c-d, S7) reveals a tripartite channel assembly (~370 kDa) having an overall structure and subunit arrangement similar to other Ca_V_1 and Ca_V_2 complexes (RMSD_Cα_ = 1.95Å, 1.71Å, and 2.37Å for Ca_V_α_1_ from Ca_v_1.1, Ca_v_1.3, and Ca_v_2.2, respectively)^40–42^ and a Ca_V_α_1_ subunit structure similar to that of single component Ca_V_3s (RMSD_Cα_ = 2.28Å for Ca_V_α_1_ from Ca_v_3.1)^43^ (Fig. S9a-d). As with other Ca_V_ structures^40–42^, the Ca_V_1.2 pore inner gate is closed and all four voltage sensors are in the ‘up’ conformation, consistent with a depolarized state and similar to those in the closely related Ca_V_1.3^41^ (Figs. S9e-h and S10a-d). Two notable features stand out as not having been previously described in other Ca_V_ structures. The first is the presence of a free leucine, a Ca_V_α_2_δ ligand^6,44,45^ and competitor of gabapentin^6,33^, in a hydrophobic pocket of the Ca_v_α_2_δ-1 Cache1 domain^46^ that is the likely binding site for gabapentinoid drugs^6,7,47^. The local resolution of this site is excellent (2.0-2.4Å) and shows the free leucine bound to a pocket lined by Trp207, Val209, Tyr219, Trp225, Tyr238, Arg243, Trp245, Tyr452, Asp454, Ala455, Leu456, and Asp493 (Figs. 2b, S7j, S10e) in good agreement with prediction of this pocket as the leucine binding site^47^. The core backbone of the bound amino acid, shared with gabapentinoids^33^, is coordinated through salt bridge and hydrogen bond interactions to its carboxylate by Tyr238, Arg243, and Trp245 and a salt bridge between the amino nitrogen and Asp493 (Figs. 2b, S10e). These interactions are in line with the established importance of Arg243 (Arg217 in some isoforms)^14,48^ and Asp493^47^ for gabapentinoid drug binding and action in pain modulation^14^ and together with prior studies establishing binding competition between leucine and gabapentin^6,33^ support the identification of this Cache1 domain as the site of gabapentinoid drug binding.

**Figure 2.**
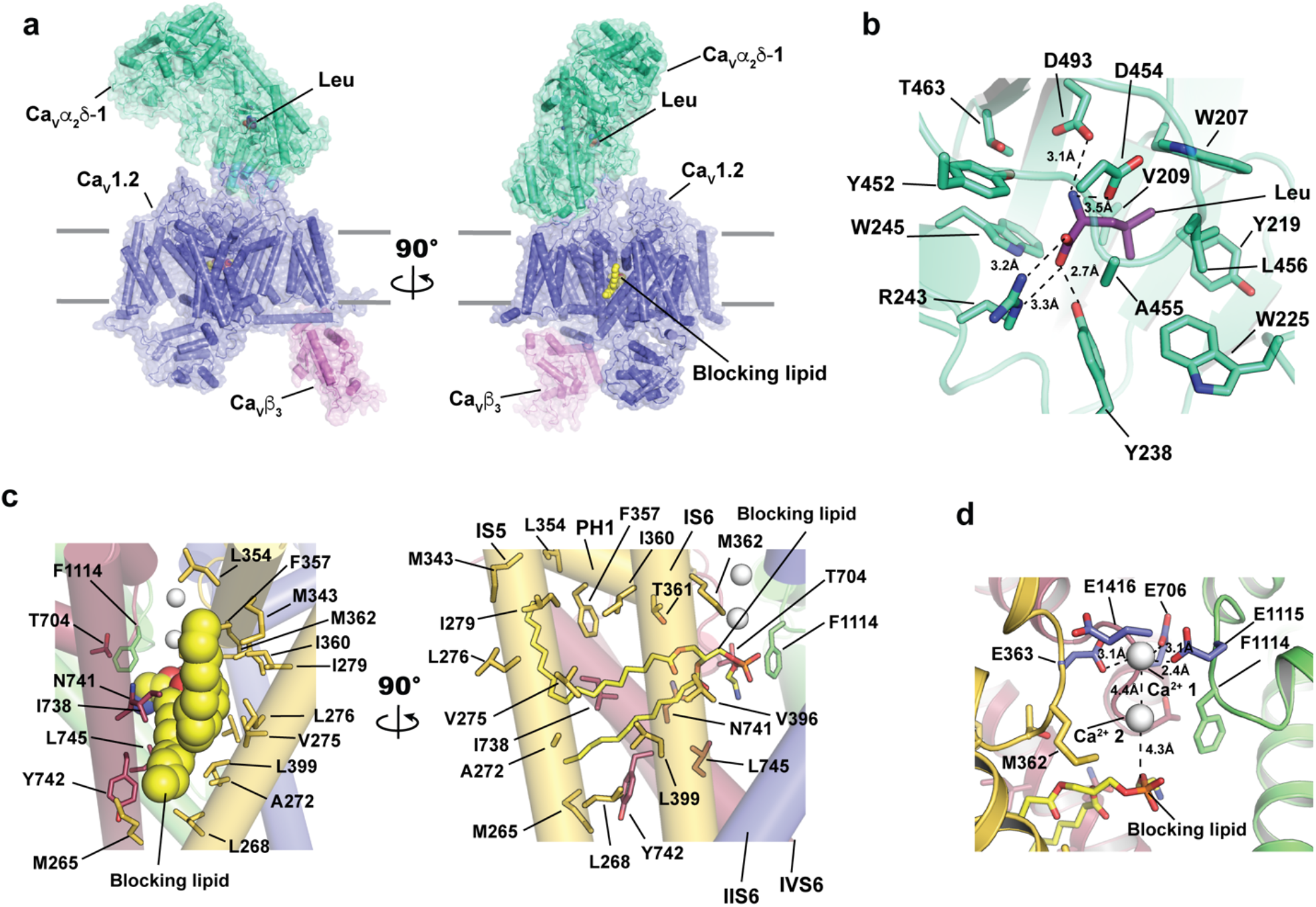
Structure of the Ca_V_1.2(ΔC)/Ca_V_β_3_/Ca_V_α_2_δ-1 channel complex. **a**, Side views of Ca_V_1.2(ΔC)/Ca_V_β_3_/Ca_V_α_2_δ-1. Subunits are colored as: Ca_V_1.2 (slate), Ca_V_β_3_ (violet), and Ca_V_α_2_δ1 (greencyan). Leucine (violet purple) and blocking lipid (yellow) are shown as space filling. Grey bars denote the membrane. **b**, Ca_V_α_2_δ-1 Cache1 ligand binding site details. Leucine (violet purple) and contacting sidechains from Ca_V_α_2_δ1 (greencyan) are shown as sticks. **c**, Blocking lipid binding site details. Ca_V_1.2 pore domains are shown as cylinders and colored. PD I (yellow orange), PD II (firebrick), PD III (lime), and PD IV (marine). Lipid contact residues are shown as sticks. Blocking lipid is shown as space filling (left) and sticks (right). **d**, Interaction of the blocking lipid and SF ions. Hallmark SF calcium binding residues are shown as sticks (slate). SF calcium ions are indicated.

The second notable feature is a lipid that we assigned as phosphatidylethanolamine (PE) (1-Heptadecanoyl-2-dodecanoyl-glycero-3-phosphoethanolamine (17:0/12:0)) that fills a hydrophobic gap between PD I and PD II (Figs. 2c and S7c). The PE hydrophilic headgroup resides in the central pore just below the selectivity filter (SF), where it coordinates the inner of two SF calcium ions (Figs. 2c-d, S7c, S10f) that match the SF ion positions in Ca_V_1.1^40,49^, Ca_V_3.1^43^, and Ca_V_3.3^50^. Similarly intrusive lipids between Ca_V_1.1 PD I and PD II^49,51^ and Ca_V_3.3 PD III and PD IV^50^ emerge into the central cavity in the presence of pore-blocking drugs. Additionally, other Ca_V_1.1^40,49,51^ and Ca_V_3.1^43^ structures identified lipid tails in the equivalent PD I/PD II interface but did not resolve the headgroup. Due to its interference with the Ca_V_1.2 ion path in the absence of drugs, we denote the Ca_V_1.2 PE as the ‘blocking lipid’. The presence of the blocking lipid polar headgroup in the central cavity in direct coordination with an SF Ca^2+^ ion shows that this lipid does not require third party molecule in the pore, such as a drug^49–51^, and raises questions about whether and how this element contributes to channel gating and pharmacology.

### Ca_V_1.2/Ca_V_β_3_ EMC holdase binding requires multiple rigid body conformational changes

Observation of the same channel complex Ca_V_1.2(ΔC)/Ca_V_β_3,_ in two different states, the EMC-bound form versus the fully assembled channel, reveals multiple, dramatic conformational changes associated with EMC binding that drastically remodel the Ca_V_ structure (Fig. 3a, Movies S2 and S3). All four of the VSDs undergo conformational changes in the EMC complex relative to the Ca_V_1.2(ΔC)/Ca_V_β_3_/Ca_v_α_2_δ-1 channel. The Ca_V_1.2 VSD I, VSD II, and VSD IV S4 segments correspond to a voltage sensor ‘up’ state in which gating charge residues (VSD I:K1, R2, R3, and R4; VSD II:R2, R3, and R4, and VSD IV:R2, R3, and R4) are on the extracellular/lumenal side of the aromatic position that defines the charge transfer center^52^ (Figs. S11a-c). Although VSD I and VSD II are similar to those in Ca_V_1.2(ΔC)/Ca_V_β_3_/Ca_v_α_2_δ-1 (Figs. S11d-e) (RMSD_Cα_ = 0.69Å and 2.25 Å, for VSD I and VSD II, respectively) both undergo substantial rigid body displacements upon EMC binding (Fig. 3a, Movies S2-S5). VSD III and VSD IV (Fig. S11f) (RMSD_Cα_ = 2.35Å for VSD IV) undergo various degrees of rearrangement (Fig. 3a, Movies S2-S3 and S6), the most dramatic of which is the unfolding of the majority of VSD III (Fig. 3a).

**Figure 3.**
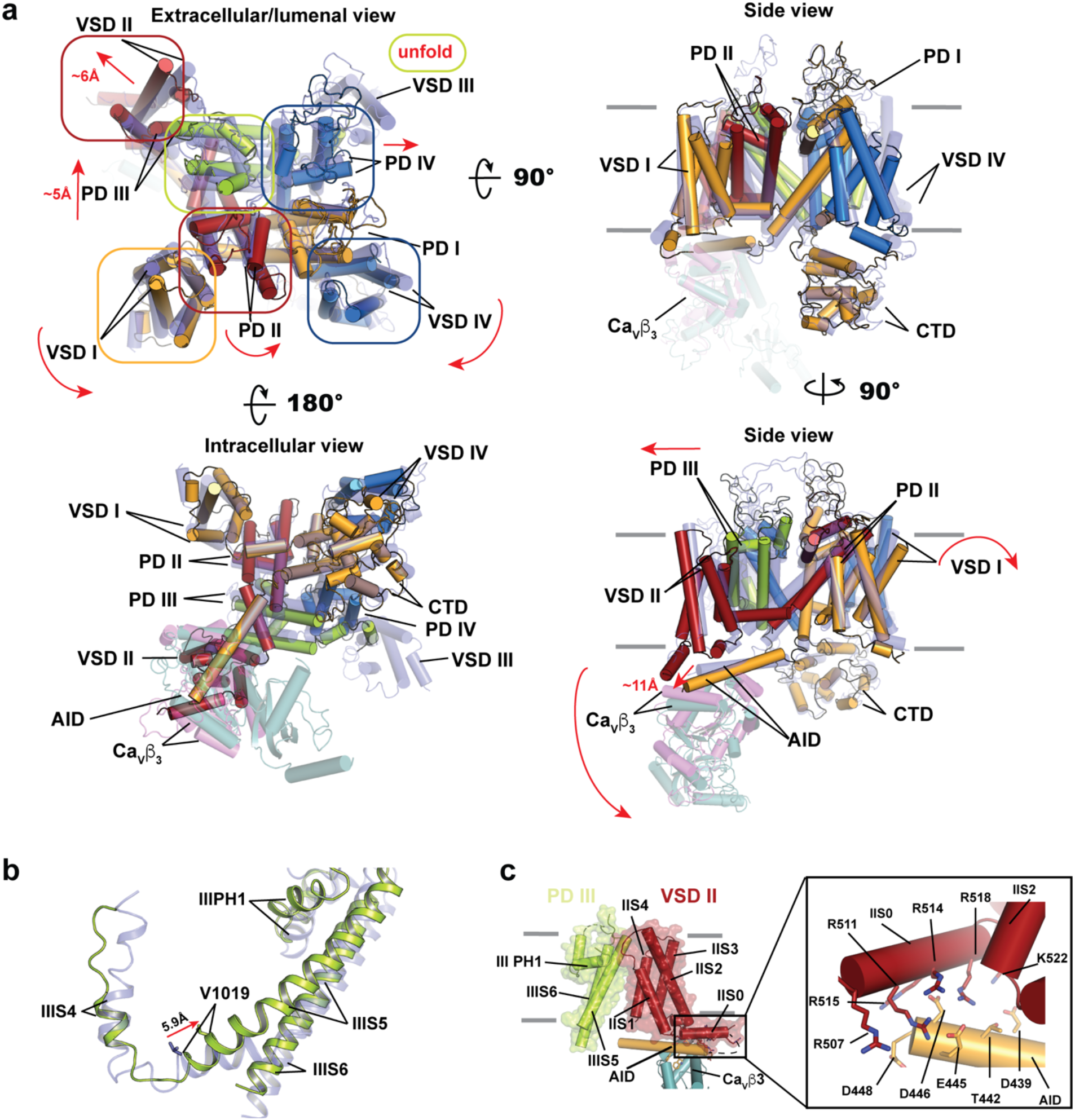
EMC interactions remodels Ca_V_ structure and extracts PD III. **a**, Superposition of Ca_V_1.2/Ca_V_β_3_ from the EMC and Ca_V_1.2/Ca_V_β_3/_Ca_V_α_2_δ-1 complexes. Ca_V_1.2 elements from the EMC complex are colored as VSD I/PD I (yellow orange), VSD II/PD II (firebrick), VSD III/PD III (lime), VSD IV/PD IV (marine). Ca_V_1.2 (slate) and Ca_V_β_3_ (violet) from Ca_V_1.2/Ca_V_β_3_/Ca_V_α_2_δ and Ca_V_β_3_ from the EMC (light teal) are semi-transparent. Red arrows indicate conformational changes between Ca_V_1.2/Ca_V_β_3_/Ca_V_α_2_δ-1 and EMC:Ca_V_1.2/Ca_V_β. Colored ovals highlight key domains reshaped by the EMC. **b**, Superposition of the S4/S4-S5 linker/PD III elements from the EMC:Ca_V_1.2/Ca_V_β_3_ complex (lime) and Ca_V_1.2/Ca_V_β3/Ca_V_α_2_δ-1 (slate). C_α_-C_α_ distance for V1019 is indicated. **c**, Interactions within the Ca_V_β:AID:VSD II:PD III subcomplex in EMC:Ca_V_1.2(ΔC)/Ca_V_β_3_. Grey bars in ‘a’ and ‘c’ denote the membrane.

Viewed from the perspective of the IS1/IS2 groove, VSD I rotates ~20° around an axis parallel with the membrane plane towards the channel periphery to engage the EMC1 TM helix (Movies S2 and S4) and rotates ~10° within the membrane plane towards VSD IV (Figs. 3a and S12a). VSD II tips away from the central pore by ~15° resulting in a displacement of the extracellular/lumenal ends of its helices by ~6Å (Figs. 3a and S12a, Movie S5). With the exception of the S4 C-terminal end, VSD III is largely disordered in the EMC complex. This lack of structure is apparent in the shape of the detergent micelle surrounding the channel (cf. Fig. S4b and d). VSD IV undergoes a complex set of changes in which S3 and S4 are repositioned relative to the S1/S2 pair, and rotates ~10° within the plane of the membrane towards VSD I (Figs. 3a and S12a, Movies S2 and S6). Thus, the relative conformations of all four VSDs change as a consequence of EMC binding.

In addition to these large VSD conformational changes, three of the four PDs in the EMC complex are also altered relative to the Ca_V_1.2(ΔC)/Ca_V_β_3_/Ca_v_α_2_δ-1 channel structure. The most striking change is PD III. This PD is partially extracted from the central pore and displaced towards the channel periphery by ~5Å (Figs. 3a and S12b, Movies S2-S3). Notably, despite some internal shifts (RMSD_Cα_ = 3.269Å), the core PD III tertiary fold comprising IIIS5, PH1, and IIIS6 helices retains a near native conformation (Fig. 12b, Movie S7), in line with the recent demonstration that VGIC PD tertiary structure is independent of quaternary structure^53^. PD III PH2 is displaced from its native position by ~15° and the large extracellular loops connected to it are disordered. The other PD tertiary structures are largely unaltered (RMSD_Cα_ = 0.91Å, 0.46Å, and 1.17Å for PD I, PD II, and PD IV, respectively), but as a consequence of PD III extraction, both PD II and PD IV undergo rigid-body changes, tipping towards and away from the central axis of the pore, respectively (Movies S2-S3, S8 and S9). PD III extraction from the central pore pulls the Domain III S4/S5 linker along the lateral side of the channel by a distance corresponding to one helical turn, ~6Å (Fig. 3c, Movie S3), disturbing VSD III and leading to its unfolding in the EMC:Ca_V_1.2(ΔC)/Ca_V_β_3_ complex (Figs. 3a and S12a).

### Ca_V_1.2/Ca_V_β_3_ binding to the EMC holdase reshapes the channel pore

Comparison of Ca_V_β_3_ in the EMC and Ca_V_1.2(ΔC)/Ca_V_β_3_/Ca_v_α_2_δ-1 complexes reveals conformational changes associated with EMC intracellular cap binding that explain the mechanism of PD III extraction from the central pore. To bind to EMC2 and EMC8, Ca_V_β_3_ rotates ~5° away from its Ca_V_1.2(ΔC)/Ca_V_β_3_/Ca_v_α_2_δ-1 position close to the membrane (Figs. 3a and S12c) and transmits this conformational change to the channel via the Ca_V_1.2 AID helix. This element remains bound to Ca_V_β_3_ and tilts ~15° away from the membrane plane, displacing its C-terminal end by ~11Å relative to its Ca_V_1.2(ΔC)/Ca_V_β_3_/Ca_v_α_2_δ-1 position (Figs. 3a and S12c). AID helix repositioning enables four AID acidic residues (Asp439, Glu445, Asp446, and Asp448) opposite to its Ca_V_β_3_ interface to interact with five VSD II IIS0 helix positively charged positions (Arg507, Arg511, Arg514, Arg515, and Arg518) (Fig. 3c). This interaction is not seen in Ca_V_1.2(ΔC)/Ca_V_β_3_/Ca_v_α_2_δ-1 (Fig. S12d) and is enabled because IIS0 becomes more helical in the EMC complex (Fig. S12d). These electrostatic interactions provide a physical link that pulls VSD II away from the channel central axis as a consequence of the AID tilt (Fig. 3a). Because VSD II IIS1 and IIS4 extracellular/lumenal ends make numerous contacts with PD III IIIS5 and PH1 (Figs. 3c and S12e) that are maintained in both structures, this VSD II movement pulls PD III away from the central pore. Hence, the PD III/VSD II/AID/Ca_V_β_3_ assembly acts as single unit whose positions are transformed due to the EMC:Ca_V_β_3_ interaction. The consequence of these changes is the partial extraction of PD III from the central pore assembly, displacement of helical S4/S5 linker between IIIS4 and IIIS5 by ~6Å (Movies S2-S3) and destabilization of VSD III (Figs. 3a-b).

PD III extraction has dramatic consequences for the central ion conduction pathway. The overall pore diameter increases due to major changes at its two narrowest points, the SF and inner gate (Figs. 4a-b,and S13). These structures widen by ~1.1Å and 1.0Å, respectively, yielding a water filled pore that bridges the channel cytoplasmic and lumenal sides (Figs. 4b, S13). SF expansion results in changes affecting the SF glutamate ring outer ion coordination site (Glu363, Glu706, Glu1115, and Glu1416)(Fig. 4c). Inner gate widening creates a pathway in which we find density corresponding to a bound glycol-diosgenin (GDN) molecule (Figs. 4d-g, S5f, and S13). This GDN sits at a position similar to that seen in a drug-bound CaV1.1/verapamil complex having a widened inner gate^49^ and may mimic a natural lipid similar to GDN such as cholesterol that would serve as a plug to prevent ion leak through the EMC-bound pore. Hence, we denote this position as ‘the lipid plug’ (Figs. 4d, f, and g). Thus, the association of the Ca_V_1.2/Ca_V_β_3_ complex with the EMC pulls and twists various channel elements, mostly through rigid body domain movements, that splay open the central pore.

**Figure 4.**
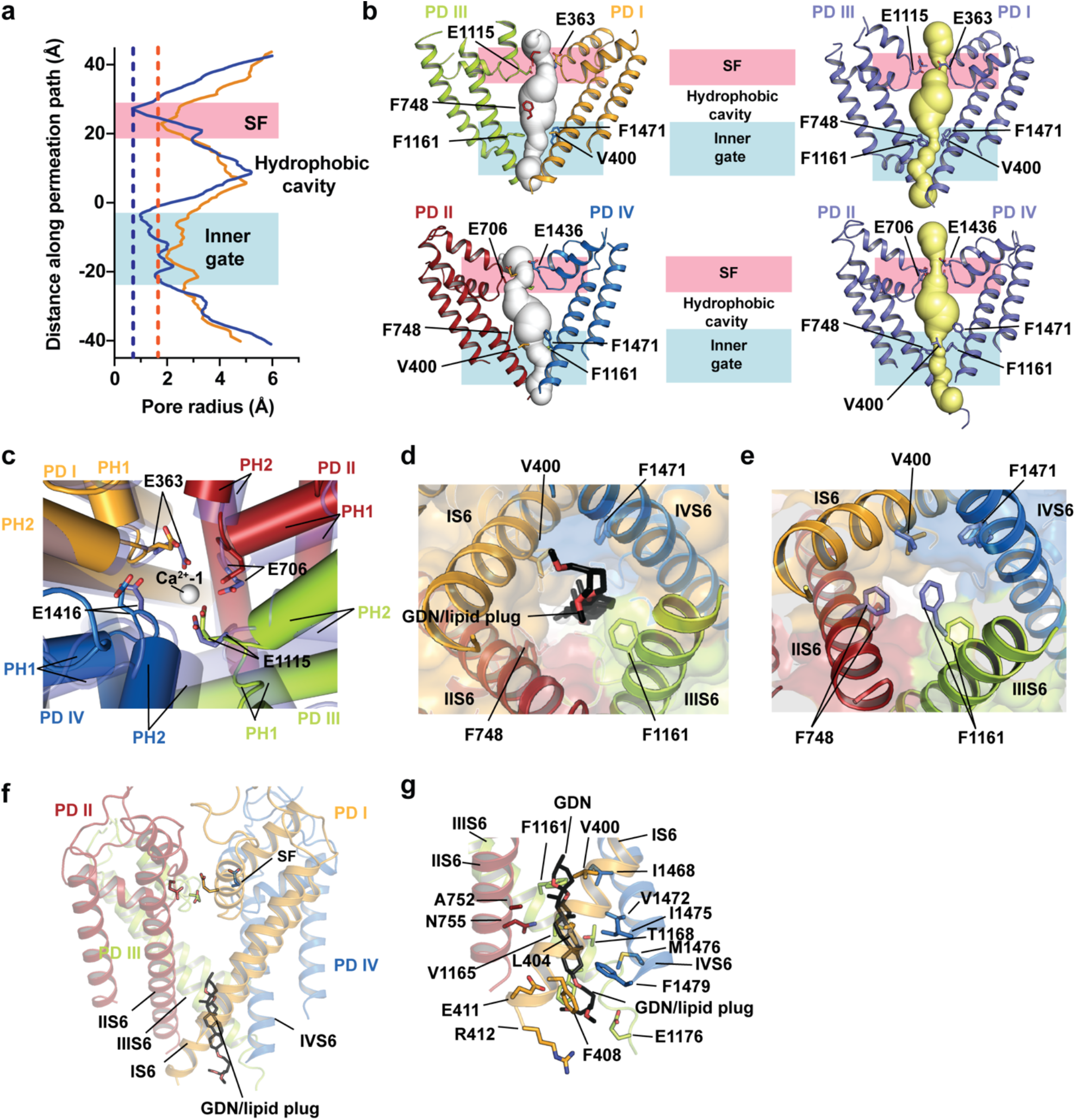
EMC association causes Ca_V_1.2 pore structural changes. **a**, Ca_V_1.2 pore profile comparison for EMC:Ca_V_1.2(ΔC)/Ca_V_β_3_ (orange) and Ca_V_1.2(ΔC)/Ca_V_β_3_/Ca_V_α_2_δ (blue), calculated with HOLE^87^. EMC: SF = 1.78 Å, inner gate = 1.98 Å; Ca_V_1.2: SF = 0.65 Å, inner gate = 0.98 Å. **b**, Side views of EMC:Ca_V_1.2(ΔC)/Ca_V_β_3_ (left) and Ca_V_1.2(ΔC)/Ca_V_β_3_/Ca_V_α_2_δ (right) pore profiles calculated with MOLE^88^. SF, central cavity, and hydrophobic gate regions are indicated (magenta, white, and blue, respectively). EMC:Ca_V_1.2(ΔC)/Ca_V_β_3_ pore domains are PD I (yellow orange), PD II (firebrick), PD III (lime), and PD IV (marine). Ca_V_1.2 pore domains are (slate). SF and intracellular gate residues are shown. **c**, Comparison of the Ca_V_1.2 SF filter regions in the EMC:Ca_V_1.2(ΔC)/Ca_V_β_3_ pore (PD I (yellow orange), PD II (firebrick), PD III (lime), and PD IV (marine)) and Ca_V_1.2(ΔC)/Ca_V_β_3_/Ca_V_α_2_ δ-1 (semitransparent, slate). Ca^2+^-1 from Ca_V_1.2(ΔC)/Ca_V_β_3_/Ca_V_α_2_δ-1 is shown as a white sphere. **d** and **e**, view of the Ca_V_1.2 intracellular gate from **d**, EMC:Ca_V_1.2(ΔC)/Ca_V_β_3_ and **e**, Ca_V_1.2(ΔC)/Ca_V_β_3_/Ca_V_α_2_δ-1. GDN is black and shown as sticks. Pore domain colors are PD I (yellow orange), PD II (firebrick), PD III (lime), and PD IV (marine). **f**, and **g**, Side views of the EMC:Ca_V_1.2(ΔC)/Ca_V_β_3_ pore showing **f**, global and **g**, detailed views. SF glutamates in ‘e’ and GDN contacting residues in ‘f’ are shown as sticks. PD elements are labeled.

### Ca_V_1.2/Ca_V_β_3_ binding to the EMC holdase raises the brace/crossbar helix to an ‘up’ position

Comparison with structures of the human EMC determined in lipid nanodiscs^26,29^ shows that Ca_V_1.2/Ca_V_β_3_ binding causes a large conformational change in the EMC lumenal domain comprising EMC1, EMC4, EMC7 and EMC10 in which the EMC top β-propeller domain moves by~11.5Å lateral to the membrane plane (Fig. 5a, S14a-b). This tilt lifts and translates the EMC1 brace/crossbar helix towards the client by ~6Å and ~10Å, respectively (Figs. 5a-b and S14a-b, Movies S10 and S11), allowing it to engage the VSD I/PD II interface (Fig. 1e) in an interaction that induces the folding of one helical turn at the brace/crossbar helix C-terminal end (residues Trp503-Tyr507) (Fig. 1e, S14a-b). We designate this as the ‘up’ conformation. Most of the remainder of the EMC complex is unchanged, with the exception of clockwise rotations of the transmembrane portions of EMC3, EMC6, and EMC5 (RMSD_Cα_ = 1.44Å, 0.91Å, and 0.63Å for EMC3, EMC5, and EMC6, respectively) and a slight counterclockwise repositioning of the EMC1 TM (RMSD_Cα_ = 0.62Å) (Fig. 5b, Movie S12). By contrast, in the detergent complex that lacks EMC7^26^, the lumenal domain and brace/crossbar helix are in an ‘up’ position similar to EMC:Ca_V_1.2(ΔC)/Ca_V_β_3_ complex (Fig. S14c) (RMSD_Cα_ = 0.84Å), even though its C-terminal helical structure is shorter than that in the EMC:Ca_V_1.2(ΔC)/Ca_V_β_3_ complex and stops at Leu502. These conformational differences suggest that brace/crossbar helix client engagement stabilizes the ‘up’ position by biasing the natural conformational landscape sampled by the apo-EMC. Up state stabilization could allow ER lumen chaperones to recognize client loaded EMCs and signal its holdase function consistent with proposals that the different conformations are associated with client-loaded states^26^. Most importantly, the structure establishes a role for the EMC1 subunit in holdase function.

**Figure 5.**
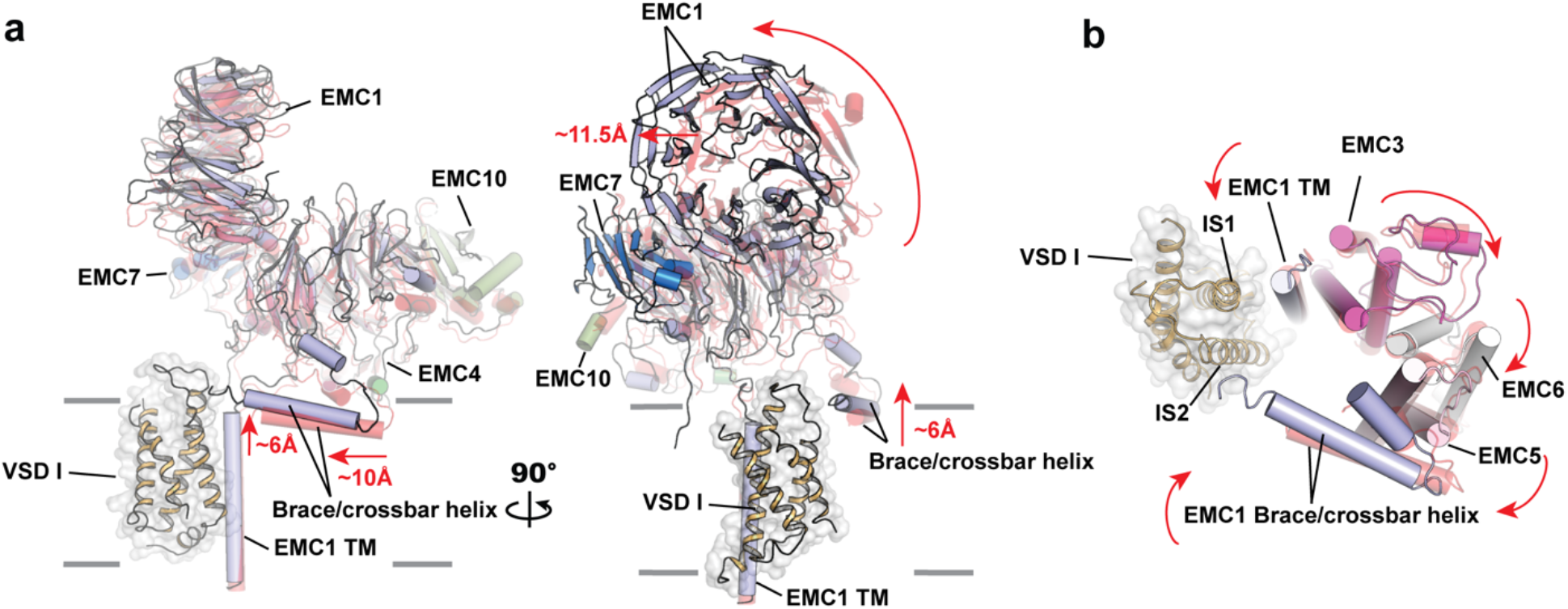
EMC structural changes between the apo-EMC and EMC:Ca_V_1.2(ΔC)/Ca_V_β_3_ client-loaded states. **a**, Side view of apo-EMC (6WW7)^29^ and EMC:Ca_V_1.2(ΔC)/Ca_V_β_3_-bound conformations showing EMC1 (light blue), EMC4 (forest), EMC7 (marine) and EMC10 (smudge) from the EMC:Ca_V_1.2(ΔC)/Ca_V_β_3_-bound state and the same subunits form apo-EMC (semi-transparent red). VSD I is shown as ribbons (yellow orange). Red arrows indicate movements of the EMC1/EMC4/EMC7/EMC10 lumenal domain assembly from the apo-to client-bound state. Grey bars denote the membrane. **b**, Lumenal view of the apo-EMC (6WW7)^29^ (semi-transparent red) and EMC:Ca_V_1.2(ΔC)/Ca_V_β_3_-bound conformations of EMC1 (light blue), EMC3 (light magenta), EMC5 (light pink), and EMC6 (white) transmembrane helices. VSD I is shown as ribbons (yellow orange). Red arrows indicate movements from the apo to client-bound states.

### EMC and Ca_V_α_2_δ binding to the Ca_V_1.2/Ca_V_β_3_ core are mutually exclusive

Superposition of the EMC:Ca_V_1.2(ΔC)/Ca_V_β_3_ complex and Ca_V_1.2/Ca_V_β_3_/Ca_V_α_2_δ-1 shows that the EMC and Ca_V_α_2_δ interact with the same Ca_V_1.2 side in a mutually exclusive manner (Fig. 6a). There are clashes with Ca_V_α_2_δ-1 all along the lumenal parts of EMC1 as well as with the EMC1 brace/crossbar helix (Figs. 6a and S15a). Notably, three EMC-bound Ca_V_1.2 elements that are displaced from their positions in Ca_V_1.2/Ca_V_β_3_/Ca_V_α_2_δ-1, VSD I, PD II, and PD III, each interact with Ca_V_α_2_δ in the fully assembled channel (Fig. S15b). Rigid body rotation of VSD I between its EMC-bound and Ca_V_α_2_δ-bound positions (Fig. 3a) moves Asp151 ~5Å to complete the coordination sphere of the calcium ion bound to the Metal Ion Dependent adhesion site (MIDAS) of the Ca_V_α_2_δ von Willebrand factor type A (VWA) domain (Fig. S15b). This VSD I residue is conserved in Ca_V_1s and Ca_V_2s (Fig. S8c) and its equivalent coordinates a similar Ca_V_α_2_δ-bound Ca^2+^ in Ca_V_1.1^40^ and Ca_V_2.2^42^. Ca_V_α_2_δ VWA MIDAS coordination of Ca^2+^ is critical for Ca_V_α_2_δ binding to Ca_V_1.2^54^ and Ca_V_2.2^55^ and is central to the ability of Ca_V_α_2_δ to promote plasma membrane channel trafficking of Ca_V_1.2, Ca_V_2.1, and Ca_V_2.2 ^54–56^. Hence, we denote this tripartite interaction as the ‘Ca^2+^ staple’ due to its importance for assembled state stabilization.

**Figure 6.**
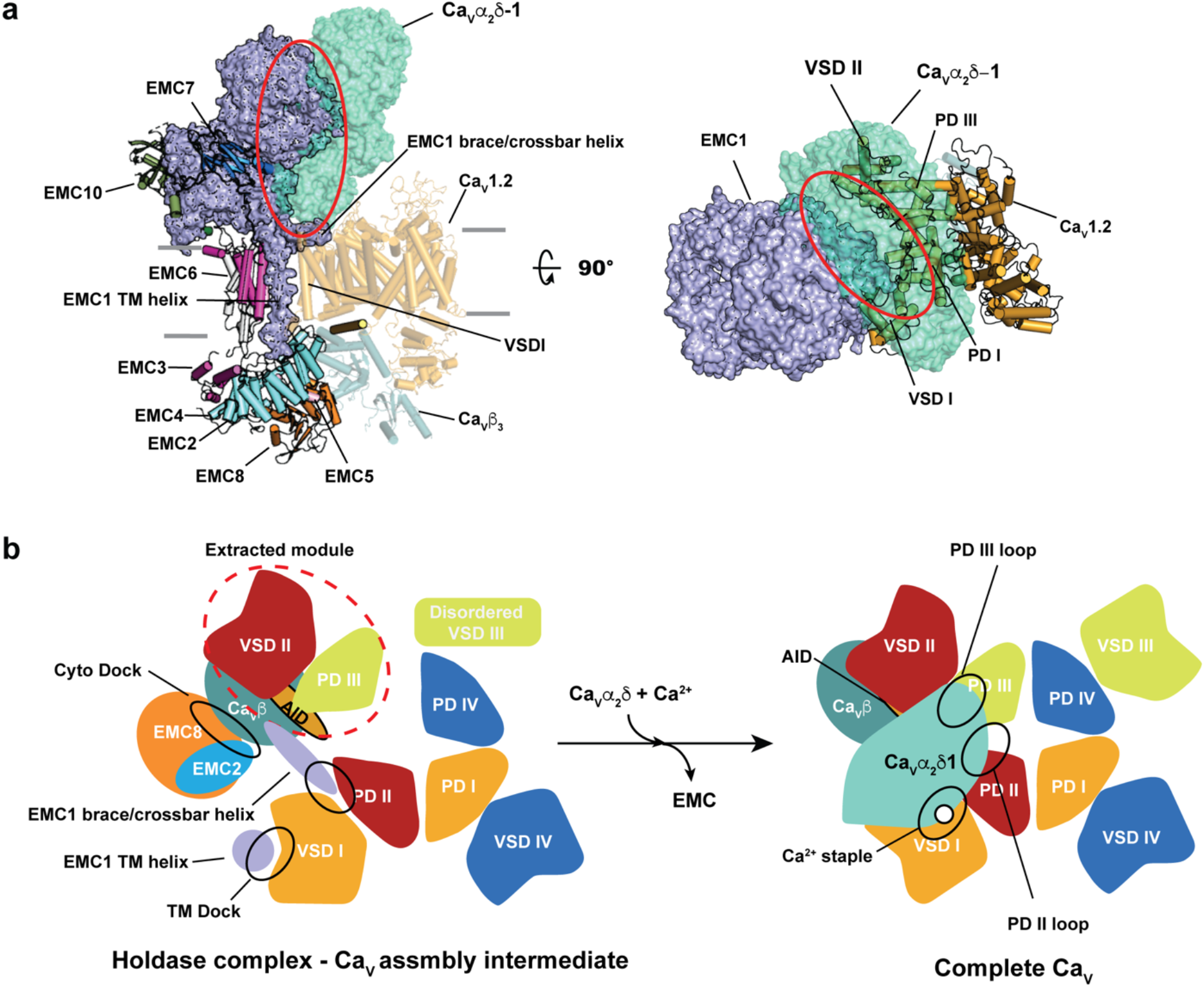
Interactions of the EMC holdase and Ca_V_α_2_δ with the core Ca_V_1.2/Ca_V_β_3_ complex are mutually exclusive. **a**, Superposition of Ca_V_α_2_δ-1 (semi-transparent, aquamarine) from the Ca_V_1.2(ΔC)/Ca_V_β_3_/Ca_V_α_2_δ-1 structure with the EMC:Ca_V_1.2/Ca_V_β_3_ complex. EMC1 surface is shown. Red oval highlights clash regions. Colors of the EMC:Ca_V_1.2/Ca_V_β_3_ complex are as in Figure 1a. Grey bars denote the membrane. **b**, Schematic showing of the conformational changes and interaction sites in the exchange between the EMC:Ca_V_/Ca_V_β holdase complex and assembled Ca_V_/Ca_V_β/Ca_V_α_2_δ channel. Black ovals indicate key interaction sites in each complex.

Movement of PD II and PD III to their Ca_V_1.2/Ca_V_β_3_/Ca_v_α_2_δ-1 positions and restoration of pore domain intersubunit interactions (Figs. 3b and 6b) appears linked to establishment of interactions with the Ca_V_α_2_δ VWA and Cache1 domains, respectively (Fig. S15b) and agrees with the important role of PD III in Ca_V_α_2_δ binding^57^. Notably, the large PD III extracellular loops linking PH1-IS5 and PH2-IS6 are disordered in the EMC:Ca_V_1.2(ΔC)/Ca_V_β_3_ complex but make extensive, structured interactions upon Ca_V_α_2_δ binding (Fig. S15b), indicating that folding of these elements is Ca_V_α_2_δ-dependent. This interaction, together with Ca^2+^ staple coordination are central to the consolidation of the native Ca_V_1.2/Ca_V_β_3_/Ca_v_α_2_δ-1 assembly and strongly suggest that the hand off of the Ca_V_1.2/Ca_V_β_3_ complex from the EMC to Ca_V_α_2_δ is a critical step in channel assembly.

### Ca_V_α_1_ associates with the EMC independently and is stabilized by Ca_V_β

The strong association of the Ca_V_1.2(ΔC)/Ca_V_β_3_ complex and EMC (Figs. S2a and S16a) prompted us to ask whether each individual channel subunit could bind the EMC. To this end, we expressed Ca_V_1.2(ΔC) or Ca_V_β_3_ and followed a purification procedure similar to the one that yielded the EMC:Ca_V_1.2(ΔC)/Ca_V_β_3_ complex to assay for EMC complex formation. Purified Ca_V_1.2(ΔC) yielded a broader SEC peak than the EMC:Ca_V_1.2(ΔC)/Ca_V_β_3_ complex (Figs. S16a-b) and contained EMC subunits identified by SDS-PAGE and mass spectrometry (Figs. S16b-c, Table S1). Quantitative comparison found lower proportions of EMC subunits relative to Ca_V_1.2(ΔC) than seen for the EMC:Ca_V_1.2(ΔC)/Ca_V_β_3_ complex (Fig. S16c), suggesting that Ca_V_1.2(ΔC) binds the EMC less tightly than the Ca_V_1.2(ΔC)/Ca_V_β_3_ pair. By contrast, Ca_V_β_3_ purification yielded no EMC subunits (Fig. S1d-e, Table S1). This sample had two lower molecular weight bands corresponding to Ca_V_β_3_ NK and SH3 domains (Fig. S16d), which are known to associate^37,58^ and likely result from Ca_V_β_3_ proteolytic cleavage. Hence, the data indicate that the Ca_V_ pore forming subunit can stably associate with the EMC and that this association is strengthened by Ca_V_β binding, in line with the observed EMC:Ca_V_1.2(ΔC)/Ca_V_β_3_ multipoint interactions. Such interactions may explain the ability of Ca_V_β subunits to protect Ca_V_1.2^59^ and Ca_V_2.2^60^ pore-forming subunits from degradation by the proteasome and the ERAD pathway^59^ in a variety of cell types, including neurons^59,60^.

## Discussion

The profound effects that the intracellular Ca_V_β^5^ and extracellular Ca_V_α_2_δ^6,7^ auxiliary subunits have on Ca_V_1 and Ca_V_2 function and trafficking have been known for more than 30 years^5,10-13,15^. Yet, apart from the role of the Ca_V_β-AID interaction in folding the short Ca_V_α_1_ AID helix^38,61^ there is scant direct structural information regarding how Ca_V_β and Ca_V_α_2_δ act in channel biogenesis and assembly. The discovery of the EMC:Ca_V_1.2(ΔC)/Ca_V_β_3_ complex and evidence that Ca_V_1.2 can bind the EMC alone offers direct insight into this process and reveals unexpected mechanisms by which the EMC acts as a client protein holdase.

Various EMC domains have been suggested to function in transmembrane segment insertion into the lipid bilayer^26,28,29,39^ or serve as holdase sites^26^. Surprisingly, none of these elements form the TM dock and Cyto dock sites that bind Ca_V_1.2(ΔC)/Ca_V_β_3_, a membrane protein complex having a different topology than EMC tail anchored protein clients^9^. Thus, our EMC:client structure expands the idea that the EMC utilizes diverse elements for different functions^26^. The EMC1 TM dock uses a lumenal side salt bridge to an EMC1 residue implicated in client binding, Asp961^30^, and a cytoplasmic side cation-π interaction^35^ with EMC1 Arg981 (Figs. S8a-b) to bind the Ca_V_1.2 VSD I (Figs. 1b-d, S8a-c). The conserved nature of this interface strongly suggests that other Ca_V_1s and Ca_V_2s interact with the EMC in a similar manner. The observation that the Ca_V_α_1_ subunit appears to bind the EMC complex on its own (Fig. S16b) suggests that EMC interactions with VSD I, the first channel transmembrane domain to be synthesized during translation^2,18^, may be an early stabilization step during biogenesis. Augmentation of the TM dock site by VSD I and PD II interactions with the EMC1 brace/crossbar helix interactions that induce the ‘up’ conformation of the EMC lumenal subassembly (EMC1/4/7/10) (Figs. 5a and S14a-b) may signal the client loaded status of the EMC to lumenal factors or other chaperones^26^. Taken together, these findings define the TM dock as a central hub of EMC holdase function.

Ca_V_β binding to the EMC Cyto dock involves EMC2 and EMC8 regions not previously known to bind client proteins and establishes the EMC intracellular domain as an important EMC:client interaction site. Ca_V_β binding to pore-forming Ca_V_α_1_ subunits not only increases the cell surface expression of Ca_V_1^59,62,63^ and Ca_V_2^64^ channels but also protects Ca_V_α_1_ subunits from degradation by the proteasome^60^ and the ERAD pathway^59^. The integral part that Ca_V_β plays in the EMC:Ca_V_1.2(ΔC)/Ca_V_β_3_ complex strongly suggests that this protection from degradation occurs though stabilization of the EMC holdase:channel complex and points to a new mechanism by which Ca_V_β binding influences the fate of Ca_V_α_1_ subunits.

The most dramatic consequence of the EMC:Ca_V_β interaction is the orchestration of the coordinated displacement of Ca_V_1.2 pore forming subunit AID, VSD II, and PD III element in a manner that results in the partial extraction of PD III from the channel pore (Figs. 3a and S12b, Movies S2 and S3). Together with the rotation of VSD I away from its native position, these changes dramatically reshape the extracellular portions of the channel adjacent to the EMC relative to their conformations in the Ca_V_1.2/Ca_V_β_3_/Ca_V_α_2_δ-1 complex. Notably, the EMC and Ca_V_α_2_δ-1 bind the same side of the Ca_V_α/Ca_V_β core in a mutually exclusive manner (Fig. 6a). Together, these conformational changes suggest that the role of the EMC is to retain the Ca_V_α/Ca_V_β complex in the ER and prepare it for Ca_V_α_2_δ binding.

Association of Ca_V_α_2_δ with the pore-forming subunit channel is governed by Ca_V_α_2_δ VWA MIDAS Ca^2+^ binding at a step thought to occur in the ER lumen^56^ and is crucial for trafficking^17,65^. In agreement with these observations, our structural data point to a central role for the ‘Ca^2+^ staple’ that bridges the Ca_V_α_2_δ VWA MIDAS domain and VSD I in the transition between the EMC-bound and Ca_V_α_2_δ-bound complexes. The structures show that handoff from the EMC:Ca_V_α/Ca_V_β complex to Ca_V_α_2_δ involves a rigid body rotation of VSD I that allows the conserved Asp151, crucial for Ca_V_α_2_δ-dependent Ca_V_1^54^ and Ca_V_2^55^ trafficking, to complete the Ca^2+^ staple coordination sphere (Figs. S8c and S15b) and stabilize interaction with Ca_V_α_2_δ. Such an exchange would permit the extracellular PH1-S5 and PH2-S6 PD Ioops that are disordered in the EMC:Ca_V_1.2/Ca_V_β_3_ complex to form extensive, structured interactions with Ca_V_α_2_δ VWA and Cache1 domains (Fig. S15b), in agreement with the importance of PD III in Ca_V_α_2_δ binding^57^. Establishing native Ca_V_α_2_δ interactions with PD III would require Ca_V_β release from the Cyto-dock and expulsion the lipid plug (Figs. 4d, f, and g) as the pore closes. Whether such steps happen in an ordered manner or in a capture-release competition between the EMC and Ca_V_β_2_δ are important unanswered questions. Taken together, our data strongly suggest that the extensive conserved interactions of Ca_V_α and Ca_V_β subunits with the EMC play a previously unrecognized role in the ability of Ca_V_α_2_δ to promote trafficking of Ca_V_1s and Ca_V_2s^10,13,66^, suggesting that this exchange is a key step in channel biogenesis and quality control and providing a new framework to study Ca_V_ biogenesis. Exchange between the EMC holdase and Ca_V_α_2_δ is likely also to be influenced by Ca_V_ domains that enhance forward trafficking^67^, Ca_V_-directed drugs^6,7,31^, especially the gabapentinoids that target Ca_V_α_2_δ^14,16,17^, and Ca_V_ subunit disease mutations^4,5,32^.

The EMC has been linked to the biogenesis of a variety of ion channels^21–23^, including one from the VGIC superfamily^22^. Given the conserved structure of VSD within the VGIC superfamily^1^, it seems likely that the EMC may participate in the biogenesis of other VGICs through TM dock interactions with this element similar to those defined here. By contrast, the binding of the Cyto dock site involves a unique subunit of Ca_V_1 and Ca_V_2 channels, Ca_V_β^5^. Interaction with this part of the EMC largely involves a Ca_V_β loop structure that could have structural equivalents in other proteins. Defining the relative roles of the TM and Cyto dock sites for binding Ca_V_s and other clients is an important line of study enabled by the structural framework presented here. Furthermore, many VGICs bear disease mutations that are not easily rationalized based on their positions in the mature channel structure. Some of these mutations may disturb interactions with the EMC that are crucial for channel assembly, maturation, and quality control.

## Supporting information

Supplementary Figs. S1-S16, Tables S1-S2, Movie legends, and References

Movie S2 Lumenal view of CaV1.2 conformational changes upon EMC binding.

Movie S3 Side view of CaV1.2 conformational changes upon EMC binding.

Movie S4 CaV1.2 VSD I conformational changes upon EMC binding.

Movie S5 CaV1.2 VSD II conformational changes upon EMC binding.

Movie S6 CaV1.2 VSD IV conformational changes upon EMC binding.

Movie S7 CaV1.2 PD III conformational changes upon EMC binding.

Movie S8 CaV1.2 PD II conformational changes upon EMC binding.

Movie S9 CaV1.2 PD IV conformational changes upon EMC binding.

Movie S10 EMC lumenal domain movement induced by client binding.

Movie S11 EMC lumenal domain movement induced by client binding.

Movie S12 EMC transmembrane domain movement induced by client binding.

## Acknowledgements

We thank T.-J. Yen and D. Bulkley for technical help and K. Brejc, H. M. Colecraft, and G. Thiel for comments on the manuscript. This work was supported by grants NIH R01 HL080050 and NIH R01 DC007664 to D.L.M., the Beckman Foundation to B.Z., the National Science Foundation GRFP DGE-2034836 to J. Montano, and American Heart Association postdoctoral fellowship to F.A.-A.. B.Z. is a Beckman Young Investigator.

## Author Contributions

Z.C. and D.L.M. conceived the study and designed the experiments. Z.C. expressed and characterized the samples. Z.C., A.M., and F.A.A. collected and analyzed cryo-EM data. J.M. and B.Z. collected and analyzed the mass spec data. Z.C. and A.M. built and refined the atomic models. Z.C. performed the functional studies. B.Z. and D.L.M. analyzed data and provided guidance and support. Z.C., A.M., F.A.-A., J.M., B.Z., and D.L.M. wrote the paper.

## Competing interests

The authors declare no competing interests.

## Data and materials availability

Coordinates and maps of the EMC:Ca_V_1.2(ΔC)/Ca_V_β_3_ complex (PDB:8EOI;EMD-28376) and Ca_V_1.2(ΔC)/Ca_V_β_3_/Ca_V_α_2_δ complex (PDB: 8EOG; EMD-28375) are deposited with the RCSB and will be released upon publication.

Mass spectrometry data is deposited in the Mass Spectrometry Interactive Virtual Environment, (massive.ucsd.edu) under identifier MSV000090434 and will be released upon publication.Requests for material to D.L.M.

## Materials and Methods

### Expression and purification of human Ca_v_1.2 and Ca_v_1.2-loaded human EMC complex

Codon-optimized cDNAs of human Ca_v_1.2 bearing a C-terminal truncation at residue 1648 (denoted Ca_V_1.2(ΔC)), a site 13 residues after the end of the IQ domain (Δ1649-2138, Uniprot Q13936-20, 1,648 residues) followed by a 3C protease cleavage site, monomeric enhanced Green Fluorescent Protein (mEGFP), and a His_8_ tag, rabbit Ca_v_α_2_δ-1 (1,105 residues, Uniprot P13806-1), and rabbit Ca_V_β_3_ (477 residues, Uniprot P54286) followed by a Strep*-*tag II sequence^68^ were synthesized (GenScript) and each subcloned into a modified pFastBac expression vector having the polyhedrin promoter replaced by a mammalian cell active CMV promoter^69^. All constructs were sequenced completely. The Strep-tag bearing rabbit Ca_V_β_3_ construct was used to generate a stable cell line in HEK293S GnTI^-^ (ATCC) using a lentiviral system as described^70^and denoted as ‘Ca_V_β_3_-stable’.

Chemically competent DH10EmBacY (Geneva Biotech) were used to generate the recombinant bacmid DNA, which was then used to transfect *Spodoptera frugiperda* (Sf9) cells to make baculoviruses for each subunit^71^. For structural studies, Ca_v_1.2 was expressed in Ca_V_β_3_-stable cells alone or together with Ca_v_α_2_δ-1 using a baculovirus transduction-based system^71^. For pulldown studies, Ca_v_1.2, Ca_V_β_3_, or both were expressed in unmodified HEK293S GnTI^-^ cells. Ca_V_β_3_-stable and HEK293S GnTI-cells were grown in suspension at 37°C supplied with 8% CO_2_ in FreeStyle 293 Expression Medium (Gibco) supplemented with 2% fetal bovine serum (FBS, Peak Serum), and were transduced with 5% (v/v) baculovirus for each target subunit when cell density reached ~2.5 × 10^6^ cells per ml. 10 mM sodium butyrate was added to cell culture 16-24 h post-transduction and the cells were subsequently grown at 30°C. Cells were harvested 48 h post-transduction by centrifugation at 5,000*g* for 30 min. The pellet was washed with Dulbecco’s phosphate buffered saline (DPBS) (Gibco) and stored at −80°C.

A cell pellet (from ~3.6 L culture) was resuspended in 200 ml of resuspension buffer containing 0.3 M sucrose, 1 mM ethylenediaminetetraacetic acid (EDTA), 10 mM Tris-HCl, pH 8.0 supplemented with 1 mM phenylmethylsulfonyl fluoride (PMSF) and 4 Pierce protease inhibitor tablets (Thermo scientific), then stirred gently on a Variomag magnetic stirrer MONO DIRECT (Thermo Scientific) at 4°C for 30 min. The membrane fraction was collected by centrifugation at 234,500*g* for 1 h and subsequently solubilized in 200 ml of solubilization buffer (buffer S) containing 500 mM NaCl, 5% glycerol (v/v), 0.5 mM CaCl2, 20 mM Tris-HCl, pH 8.0, and supplemented with 1% (w/v) glycol-diosgenin (GDN) and rotated on an Orbitron rotator II (speed mode S) (Boekel Scientific) at 4°C for 2 h. The supernatant, collected by centrifugation at 234,500*g* for 1 h, was diluted with an equal volume of buffer S to a final concentration of 0.5% GDN and incubated with anti-GFP nanobody Sepharose resin^72^at 4°C overnight. The resin was loaded on an Econo-Column chromatography column (BioRad) and then was then washed stepwise with 20 column volumes (CV) of buffer S supplemented with 0.1% (w/v) GDN, 20 CV of buffer S supplemented with 0.02% (w/v) GDN, and 20 CV of elution buffer (buffer E) containing 150 mM NaCl, and 0.5 mM CaCl_2_, 0.02% (w/v) GDN 20 mM Tris-HCl pH 8.0. The protein was eluted with 3C protease^73^ and subsequently incubated at 4°C for 2 h with 4 mL of *Strep*-tactin Superflow Plus beads (Qiagen) pre-equilibrated with buffer E. The beads were washed with 20 column volumes of buffer E and the protein was eluted with buffer E supplemented with 2.5 mM desthiobiotin. The eluent was concentrated using an Amicon Ultra-15 100-kDa cut-off centrifugal filter unit (Merck Millipore) before purification on a Superose 6 Increase 10/300 GL gel filtration column (GE Healthcare) pre-equilibrated in buffer E. Peak fractions were pooled for mass spectrometric analysis and concentrated with an Amicon Ultra – 0.5mL 100-kDa cut-off centrifugal filter unit (Merck Millipore) for cryo-EM sample preparation (1.7 mg ml^-1^ Ca_v_1.2(ΔC)/Ca_v_β_3_ sample or 2.7 mg/ml Ca_v_1.2(DC)/Ca_v_β_3_/Ca_v_α_2_δ sample).

### Two-electrode voltage clamp electrophysiology

The C-terminus truncated human Ca_v_1.2 (Δ1649-2138, designated as ‘Ca_v_1.2(ΔC)’) construct was obtained from a full-length human Ca_v_1.2 α1 (Uniprot Q13936-20) template in pcDNA3.1 (Invitrogen) using a Gibson Assembly kit (NEB). Full-length Ca_v_1.2 or Ca_v_1.2(ΔC) was co-expressed with rabbit Ca_V_β_3_ (Uniprot P54286) in pcDNA3.1 for *Xenopus* oocyte two-electrode voltage-clamp experiments. Five micrograms of Ca_v_1.2 or Ca_v_1.2(ΔC) and Ca_V_β_3_ cDNA were linearized by 30 units of restriction enzymes XhoI and NotI (NEB), respectively, at 37°C overnight. The linearized cDNA was used as the template to synthesize capped mRNA using a T7 mMessenger kit (Ambion). 100 nL of Ca_v_1.2 or Ca_v_1.2(ΔC) and Ca_V_β_3_ mixed at a 1:1 molar ratio was injected into *Xenopus* oocytes 1-2 days before recording. Two-electrode voltage-clamp experiments were performed as previously described^74^. Briefly, oocytes were injected with 50 nL of 100 mM BAPTA 2-4 minutes before recording, to minimize calcium-activated chloride currents^74,75^. For recording of Ca^2+^ or Ba^2+^ currents, bath solutions contained 50 mM NaOH, 40 mM Ca(NO_3_)_2_ or 40 mM Ba(OH)_2_, respectively, 1 mM KOH, 10 mM HEPES, adjusted to pH 7.4 with HNO_3_. Electrodes were filled with 3 M KCl and had resistances of 0.3–1.0 MΩ. Recordings were conducted at room temperature from a holding potential of −90 mV. Leak currents were subtracted using a P/4 protocol.

### Data quantification and statistical analysis

All the details of data analysis and statistical analysis can be found in the Method Details and figure legends. All results are from at least two independent oocyte batches.. Data were analyzed with Clampfit 10.6 (Axon Instruments). Activation curves were obtained by fitting the data with the following Boltzmann equation: Y = Y_min_ + (Y_max_ - Y_min_)/(1 + exp ((V_1/2_ - V_m_)/k), where V_1/2_ is the midpoint of activation, k is the slope factor of the activation curve, Y = G/G_max_ and Y_max_ and Y_min_ are the maximum and minimum values of G/G_max_, respectively, where G = I/(V_m_ - E_rev_) and G_max_ is the maximal macroscopic conductance, where I is the measured peak current at each test potential (V_m_). All data values are presented as mean ± SEM. ‘n’ represents the number of oocytes. Statistical significance of the observed effects was assessed by unpaired t test, using GraphPad Prism version 9.3.1 software. p < 0.01 was considered significant, unless otherwise stated.

### Stain-free gel Image analysis and in-gel digestions for mass spectrometry analysis

Purified protein samples were diluted in 4x Laemmli Sample Buffer (BioRad) containing 10% β-mercaptoethanol. Samples (1 mg of protein per well) were loaded onto a protein gel (BioRad, 4-15% Criterion TGX Stain-Free Precast Gel). Gels were run at 200V for 42 minutes and imaged on a gel imager (BioRad, ChemiDoc MP) using the ‘stain-free gel’ setting. Gels were stained with colloidal blue stain (Invitrogen) overnight at room temperature with gentle agitation. Excess stain was removed by briefly (3 min) by rinsing the gel in de-stain solution (50% water, 40% methanol, 10% acetic acid) followed by 3 × 15 min washes MilliQ water. Gel lanes were cut into thirds. Each gel slice was subsequently diced into approximately 1 mm cubes. Gel cubes corresponding to a single gel slice were transferred to a 1.5 mL microcentrifuge tube (Protein Lo-Bind, Eppendorf). 100 mM Ammonium Bicarbonate (ABC) was added to each tube (75 μL, or enough to completely cover gel cubes), and samples were incubated for 10 minutes at room temperature. ABC solution was pipetted away with gel-loading tips (VWR). 100 μL of 100 mM ABC and 10 μL of DTT (50 mM) were added to each sample before incubating at 55°C (400 rpm, for 30 minutes). Excess buffer was removed with a gel-loading tip and replaced with 100 μL of 100 mM ABC and 10 μL of 150 mM iodoacetamide. Samples were incubated in the dark for 30 minutes at room temperature. Buffer was removed, and gel cubes were washed 2x with 150 μL of 1:1 100 mM ABC:Acetonitrile on a rotator at room temperature for 10 min. After the last wash, gel cubes were dehydrated with 100 μL of acetonitrile. Excess acetonitrile was removed on a speed-vac for 30 minutes. 150 μL of 3.3 ng/μL trypsin (Promega) in ABC (50 mM) was added to each sample (0.5μug trypsin/sample) and incubated on a shaker (300 rpm) at 37°C for 16 hours.

After digestion, samples were acidified and peptides eluted through the addition of 250 μL of a 66% acetonitrile, 33% ABC (100 mM), 1% formic acid (FA) solution. Samples were then centrifuged at 10,000 xg for 2 min, and the resulting supernatants were transferred to new Protein LoBind eppi tubes (Eppendorf). Peptide elution step was repeated and corresponding supernatants were combined. Samples were then dried on a speed-vac for 2 hours.

Crude dried peptides were cleaned using ZipTip C18 columns (Millipore) as per the manufacturer’s instructions. Briefly, peptides were resuspended in 20 μL of resuspension buffer (5:95 ACN:MilliQ H_2_O, 0.1% TFA) and solubilized in a bath sonicator (VWR, 35 kHz, 10 min). A ZipTip was placed on a P10 pipettor set to 10 μL, and the C18 column was activated 2x with 10 μL of hydration buffer (50:50 ACN: MilliQ H_2_O, 0.1% TFA), followed by a single wash step with 10 μL of wash buffer (0.1% TFA in water). Peptides were loaded onto the C18 ZipTip column by taking up 10 μL of sample and slowly pipetting up and down 10x. This loading step was repeated once more before washing 2x with 10 μL of wash buffer. Finally, 10 μL of elution buffer (60:40 ACN:MilliQ H_2_O, 0.1% TFA) was taken up into the ZipTip and expelled into a fresh LoBind tube, followed by pipetting up and down 10x inside the tube. The elution step was repeated once. Eluted peptides were dried on a speed-vac for 20 min and then stored at −20°C until mass spectrometry (MS) analysis.

Before MS analysis, peptides were resuspended in loading buffer (2% ACN, 0.1% FA) and completely solubilized in a bath sonicator (VWR, 35 kHz, 10 min).

### Mass spectrometry data acquisition and analysis

#### Mass spectrometry analysis – liquid chromatography and timsTOF Pro

A nanoElute (Bruker) was attached in line to a timsTOF Pro equipped with a CaptiveSpray Source (Bruker, Hamburg, Germany). Chromatography was conducted at 40°C through a 25cm reversed-phase C18 column (PepSep) at a constant flow-rate of 0.5 μL/min. Mobile phase A was 98/2/0.1% LC/MS grade H_2_O/ACN/FA (v/v/v) and phase B was LC/MS grade ACN with 0.1% FA (v/v). During a 108 min method, peptides were separated by a 3-step linear gradient (5% to 30% B over 90 min, 30% to 35% B over 10 min, 35% to 95% B over 4 min) followed by a 4 min isocratic flush at 95% for 4 min before washing and a return to low organic conditions. Experiments were run as data-dependent acquisitions with ion mobility activated in PASEF mode. MS and MS/MS spectra were collected with *m*/*z 1*00 to 1700 and ions with *z* = +1 were excluded.

Raw data files were searched using PEAKS Online Xpro 1.6 (Bioinformatics Solutions Inc.). The precursor mass error tolerance and fragment mass error tolerance were set to 20 PPM and 0.05 respectively. The trypsin digest mode was set to semi-specific and missed cleavages was set to The human Swiss-Prot reviewed (canonical) database (downloaded from UniProt) and the common repository of adventitious proteins (cRAP, downloaded from The Global Proteome Machine Organization) totaling 20,487 entries were used. Carbamidomethylation was selected as a fixed modification. Acetylation (N-term), Deamidation (NQ) and Oxidation (M) were selected as variable modifications. The maximum number of variable PTMs per peptide was to 3.

All experiments were repeated in triplicate and combined datasets subjected to the following filtration criteria:

1. All proteins from common contaminants search database and all keratins were removed.
2. Required a protein to be found in all 3 replicates of at least one condition.
3. At least 20% coverage
4. At least 4 peptides and 4 unique peptides
5. Median Area of at least 25000 across all 3 replicates in the Ca_v_1.2(ΔC)/Ca_v_β_3_ sample (denoted as Alpha+Beta in Table S1).

Raw data files and searched datasets are available on the Mass Spectrometry Interactive Virtual Environment, a member of the Proteome Xchange consortium (massive.ucsd.edu) under the identifier MSV000090434. The filtered dataset is also available in Supplementary Table S1.

### Sample preparation and cryo-EM data acquisition

For cryo-EM, 3.5 μl of 1.7 mg ml^-1^ Ca_v_1.2(ΔC)/Ca_v_β_3_ sample or 2.7 mg/ml Ca_v_1.2(ΔC)/Ca_v_β_3_/Ca_v_α_2_δ sample was applied to Quantifoil R1.2/1.3 300 mesh Au holey-carbon grids, blotted for 4-6 s at 4°C and 100% humidity using a FEI Vitrobot Mark IV (Thermo Fisher Scientific), and plunge frozen in liquid ethane. Cryo-EM grids were screened on a FEI Talos Arctica cryo-TEM (Thermo Fisher Scientific) (at University of California, San Francisco (UCSF) EM Facility and SLAC National Accelerator Laboratory) operated at 200 kV and equipped with a K3 direct detector camera (Gatan), and then imaged on a 300 kV FEI Titan Krios microscope (Thermo Fisher Scientific) with a K3 direct detector camera (Gatan) (UCSF). Cryo-EM datasets were collected in super-resolution counting mode with a super-resolution pixel size of 0.4233 Å (physical pixel size of 0.8466 Å) using SerialEM^76^. Images were recorded with a 2.024 s exposure over 81 frames with a dose rate of 0.57 e^−^ Å^−2^ per frame. The defocus range was set from −0.9 μm to −1.7 μm.

### Imaging processing and 3D reconstruction

A total of 14,341 and 13,704 movies were collected for Ca_v_1.2(ΔC)/Ca_v_β_3_ and Ca_v_1.2(ΔC)/Ca_v_β_3_/Ca_v_α_2_δ samples, respectively. Initial image processing was carried out in cryoSPARC-3.2^77^. Raw movies were corrected for motion and Fourier binned by a factor of two (final pixel size of 0.8466 Å) with the patch motion job. Contrast transfer function (CTF) parameters of the resulting micrographs were estimated with the patch CTF job in cryoSPARC-3.2. Particles were picked by blob picking, extracted using a box size of 440 (Fourier crop box size of 220), and 2D-classified using a mask diameter of 260 Å. Particles were subsequently template picked with 2D class averages low-pass filtered to 20 Å. The resulting particles were extracted using a box size of 440 (2x Fourier cropping) and applied to one round of *ab initio* and heterogeneous refinement with C1 symmetry. Particles having reasonable 3D reconstructions (as judged by the Fourier shell correlation (FSC) curve shapes) were re-extracted without Fourier cropping and subjected to iterative rounds of *ab initio* and heterogeneous refinements. Non-uniform refinements were performed to archive 3D reconstructions with the highest resolution.

To improve the features of each 3D reconstruction, multibody refinement was carried out in RELION-3.1^78^. For Ca_V_1.2/Ca_V_β_3_, 239,729 refined particles were exported from cryoSPARC-3.2 to RELION-3.1 using the csparc2star.py (UCSF pyem v0.5. Zenodo) suite of conversion scripts (https://doi.org/10.5281/zenodo.3576630). Following a 3D refinement in RELION-3.1 using the refined map from cryoSPARC-3.2 and the exported particles, an overall 3.4 Å EM density map was obtained (EMC:Ca_v_1.2/Ca_v_β3 (ECAB) Map 1) (see Supplementary Fig. S2i for processing flow charts). To improve the map features of the flexible regions, Multibody refinement was performed in RELION-3.1 using two bodies: (i) body I containing the lumenal domain (i.e. EMC1, EMC4, EMC7, and EMC10) and the transmembrane region (Ca_V_1.2, EMC1 TM1, EMC3 TM1-3, EMC5 TM1-2, and EMC6) of the ECAB Map 1), and (ii) body II containing the cytoplasmic domain (Ca_V_β_3_, EMC2, EMC8, and the cytoplasmic segments of EMC3 and EMC5) and an overlapping portion of the transmembrane region. phenix.combine_focused_maps program was used to combine the two segments with improved features from Multibody refinement^79^. A similar strategy was applied to 269,950 particles of ECAB Map 2 to obtain a 3.6 Å consensus map and followed by feature improvement using Multibody refinement in RELION-3.1. ECAB Map 1 and ECAB Map 2 (cross correlation = 0.9836) were merged using Phenix to obtain ECAB Map 3 with best features^79^. The pool of 269,802 particles of Ca_v_1.2/Ca_v_β3/Ca_v_α2δ (CABAD) Map 1 was exported from cryoSPARC using the strategy as mentioned above and subjected to 3D refinement in RELION-3.1, yielding a 3.3 Å consensus map. This map and particle set was further used to perform a two-body Multibody refinement following the procedure mentioned above to obtain CABAD Map 2.

### Model building and refinement

ECAB Map 3 and CABAD Map 2 were used to build the model for the EMC:Ca_v_1.2(ΔC)/Ca_v_β_3_ complex and the Ca_v_1.2(ΔC)/Ca_v_β_3_/Ca_v_α_2_δ-1 complex, respectively. For the EMC:Ca_v_1.2(ΔC)/Ca_v_β_3_ complex, a preliminary model was built using the previously reported apo-EMC structure (PDB:6WW7)^29^ and the Ca_v_β_3_ subunit in the Ca_v_2.2 structure (PDB:7MIY)^42^. For the Ca_v_1.2(ΔC)/Ca_v_β_3_/ Ca_v_α_2_δ-1 complex, the Ca_v_2.2 structure (PDB:7MIY)^42^ was used as starting model. Phyre2 server^80^ was used to predict 10 models of Ca_v_1.2 using amino acid sequences (Uniprot code Q13936-20) and among them the best model was selected by comparing them with the density map. phenix.dock_in_map program was used to dock the models in respective maps^79^. Docked model and maps were further manually checked and fitted in COOT^81^. Iterative structure refinement and model building were performed using phenix.real_space_refine program^79^. The local resolution range for Ca_v_1.2 was 2.0 – 4.8 Å (Extended Data Figure S3e), which allowed model building by putting more emphasis on helix topology and large aromatic side-chain densities. However, model building was strictly followed as per the available electron density. Restraint files necessary for refinement were generated using phenix.elbow^79,82^. Final statistics of 3D reconstruction and model refinement can be found in (Table S1). The per-residue B-factors, after final refinement against the overall map, were rendered on the refined model and presented in (Figs. S3c-f). The final models were evaluated using MolProbity^83^. All figures and movies were generated using ChimeraX^84^ and Pymol package (http://www.pymol.org/pymol). Close-contact interaction analysis were performed using LIGPLOT and DIMPLOT^85,86^.

## Supplementary material

Supplementary material contains Figures S1-S16, Movies S1-S12, Tables S1-S2, and references.

## Notes

### Competing Interest Statement

The authors have declared no competing interest.

