## Supplementary Figs. S1-S16, Tables S1-S2, Movie legends, and References for "EMC holdase:Ca_V_1.2/Ca_V_β_3_ complex and Ca_V_1.2 channel structures reveal Ca_V_ assembly and drug binding mechanisms"

3 October 2022

[6]Molecular Biophysics and Integrated Bio-imaging Division
Lawrence Berkeley National Laboratory, Berkeley, CA 94720 USA

**Keywords:** EMC, voltage-gated calcium channel, holdase, $Ca_V\alpha_2\delta$, drug binding

### Figure S1

Chen *et al.*

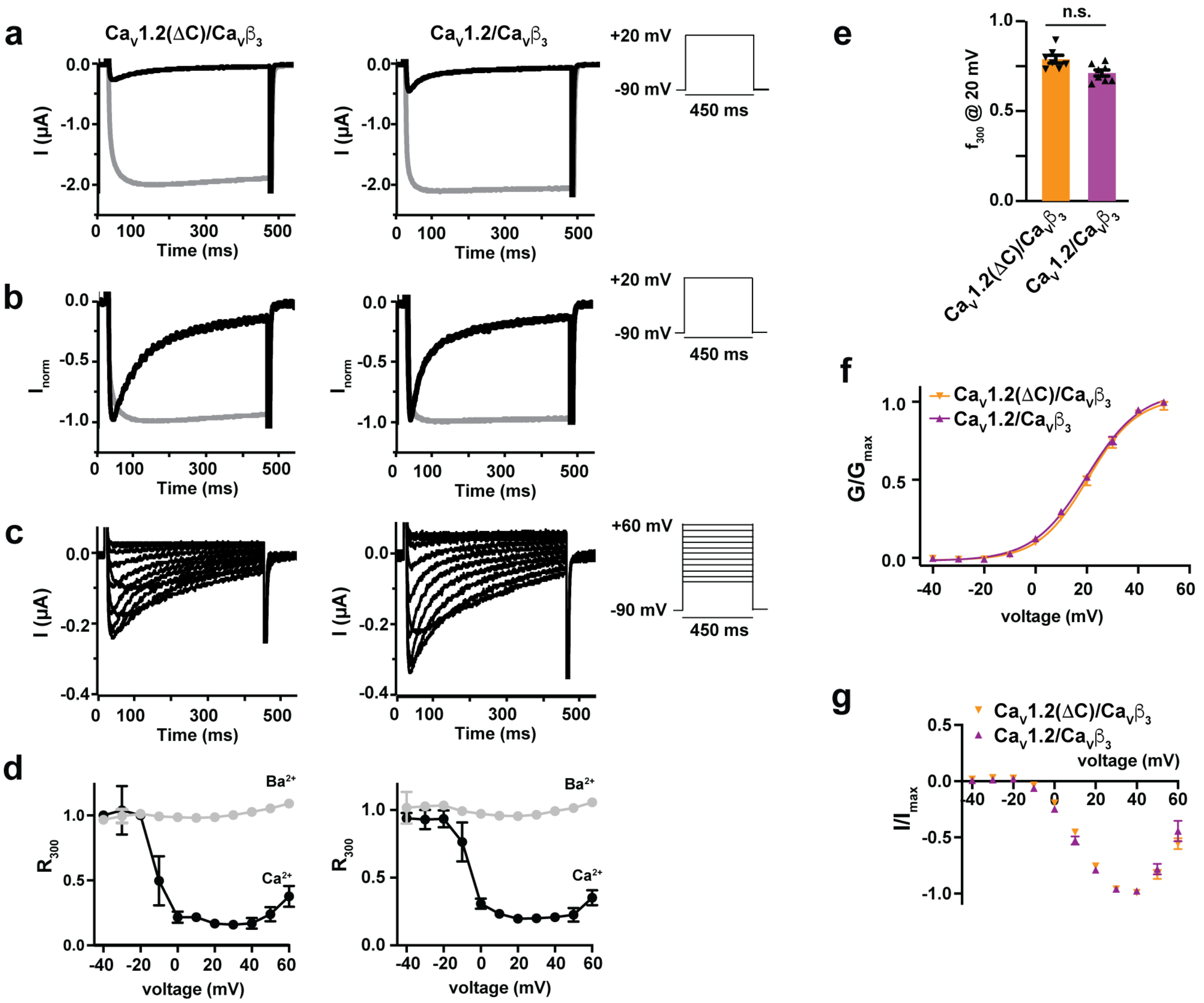


**Figure S1 $Ca_V1.2(\Delta C)/Ca_V\beta_3$ functional properties are similar to $Ca_V1.2/Ca_V\beta_3$ a**, Exemplar recordings in $Ca^{2+}$ (black) or $Ba^{2+}$ (grey) at +20 mV from *Xenopus* oocytes expressing $Ca_V1.2(\Delta C)/Ca_V\beta_3$ or $Ca_V1.2/Ca_V\beta_3$. **b**, Exemplar normalized recordings from '**a**'. **c,** Exemplar $Ca^{2+}$ currents evoked using a multi-step activation protocol with a voltage ramp from -40 mV to +60 mV. Insets in **a-c,** show protocols. **d** Fractional $Ca^{2+}$ (black) or $Ba^{2+}$ (grey) current remaining 300 ms post-depolarization ($R_{300}$) as a function of the membrane potential for channels in '**c**'. $R_{300} = I_{300}/I_0$ where $I_{300}$ and $I_0$ are current amplitudes at 300 ms post-depolarization and at the peak current, respectively. **e,** Average fraction CDI ($f_{300}$) at +20 mV, where $f_{300}$ is the difference between $Ca^{2+}$ and $Ba^{2+}$ at $R_{300}$. n.s., not significant, p>0.01. **f,** Voltage-dependent activation curves for $Ca_V1.2(\Delta C)/Ca_V\beta_3$ (orange triangles) and $Ca_V1.2/Ca_V\beta_3$ (purple triangles) channels. $V_{1/2}$ is the midpoint of activation. k is the slope factor of the activation curve. $V_{1/2}$, k and G were determined using $Ca^{2+}$ as the charge carrier and $G_{max}$ is the maximal macroscopic conductance.

$Ca_V1.2(\Delta C)/Ca_V\beta_3$: $V_{1/2}$ = 21.0 ± 1.6 mV, k = 9.8 ± 0.6. $Ca_V1.2/Ca_V\beta_3$: $V_{1/2}$ = 20.2 ± 1.0 mV, k = 10.5 ± 0.3. **g,** I-V relationships for $Ca_V1.2(\Delta C)/Ca_V\beta_3$ and $Ca_V1.2/Ca_V\beta_3$ and channels. Symbols are the same as **f**. $Ca_V1.2(\Delta C)/Ca_V\beta_3$: $E_{rev}$ = 85.80 ± 3.55 mV. $Ca_V1.2/Ca_V\beta_3$: $E_{rev}$ = 85.78 ± 5.98 mV. n = 7-8 oocytes. Data are expressed as mean values ± SEM.

**Figure S2** **Chen *et al.***

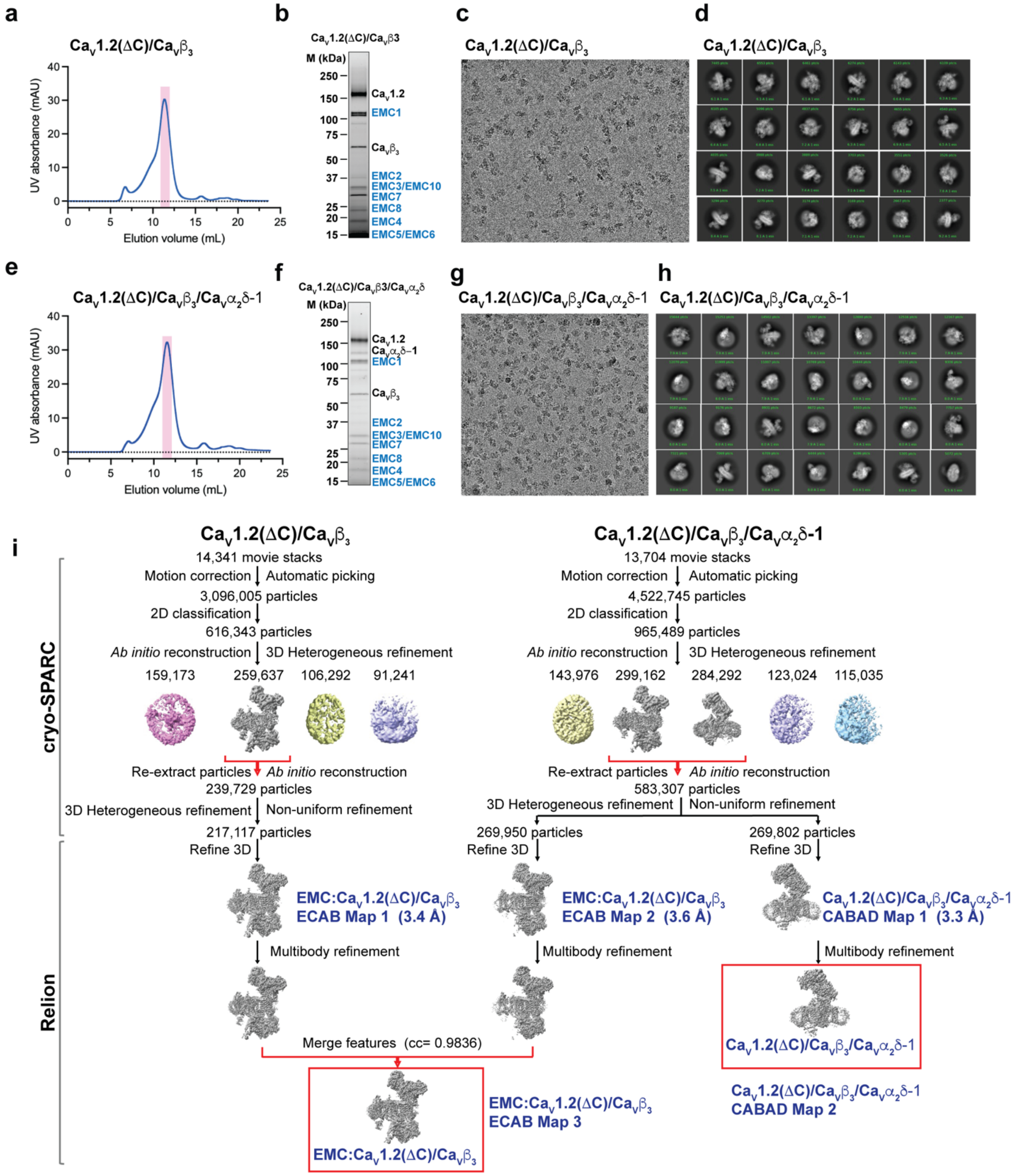


**Figure S2 EMC:CaV1.2(ΔC)/CaVβ3 and CaV1.2(ΔC)/CaVβ3/CaVα2δ-1 Cryo-EM analysis** **a-d,** Exemplars of purified CaV1.2(ΔC)/CaVβ3: **a**, SEC (Superose 6 Increase 10/300 GL). **b,** peak fraction SDS-PAGE **c,** electron micrograph (~105,000x magnification), and **d,** 2D class averages. **e-h** Exemplars of purified CaV1.2(ΔC)/CaVβ3/CaVα2δ-1: **e**, SEC (Superose 6 Increase 10/300 GL).

**f,** peak fraction SDS-PAGE **g,** electron micrographs (~105,000x magnification), and **h,** 2D class averages. **i**, Workflow for electron microscopy data processing for the $Ca_V1.2(\Delta C)/Ca_V\beta_3$ and $Ca_V1.2(\Delta C)/Ca_V\beta_3/Ca_V\alpha_2\delta$-1 samples. Initial cryoSPARC-3.2 *Ab initio* reconstruction identified a population of particles containing the EMC:$Ca_V1.2(\Delta C)/Ca_V\beta_3$ complex in the $Ca_V1.2(\Delta C)/Ca_V\beta_3$ sample and populations of particles containing either the EMC:$Ca_V1.2(\Delta C)/Ca_V\beta_3$ complex or $Ca_V1.2(\Delta C)/Ca_V\beta_3/Ca_V\alpha_2\delta$-1complex in the $Ca_V1.2(\Delta C)/Ca_V\beta_3/Ca_V\alpha_2\delta$-1 sample. Red arrows indicate the three classes that were re-extracted, subjected to multiple rounds of 3D heterogeneous classification, and exported from cryoSPARC-3.2 for further 3D refinement in RELION-3.1. This resulted in two maps for the EMC:$Ca_V1.2(\Delta C)/Ca_V\beta_3$ complex (ECAB Maps 1 and 2) and one for the $Ca_V1.2(\Delta C)/Ca_V\beta_3/Ca_V\alpha_2\delta$-1 complex (CABAD Map 1). Multibody refinement was performed in RELION-3.1 to improve the features of flexible regions of the three maps. This resulted in the final map for the $Ca_V1.2(\Delta C)/Ca_V\beta_3/Ca_V\alpha_2\delta$-1 complex (CABAD Map 2). ECAB Maps 1-2 with improved flexible features were merged (cross correlation = 0.9836) to obtain the final map for the EMC:$Ca_V1.2(\Delta C)/Ca_V\beta_3$ complex (ECAB Map 3). Red boxes indicate the final maps used for model building.

**Figure S3** **Chen *et al.***

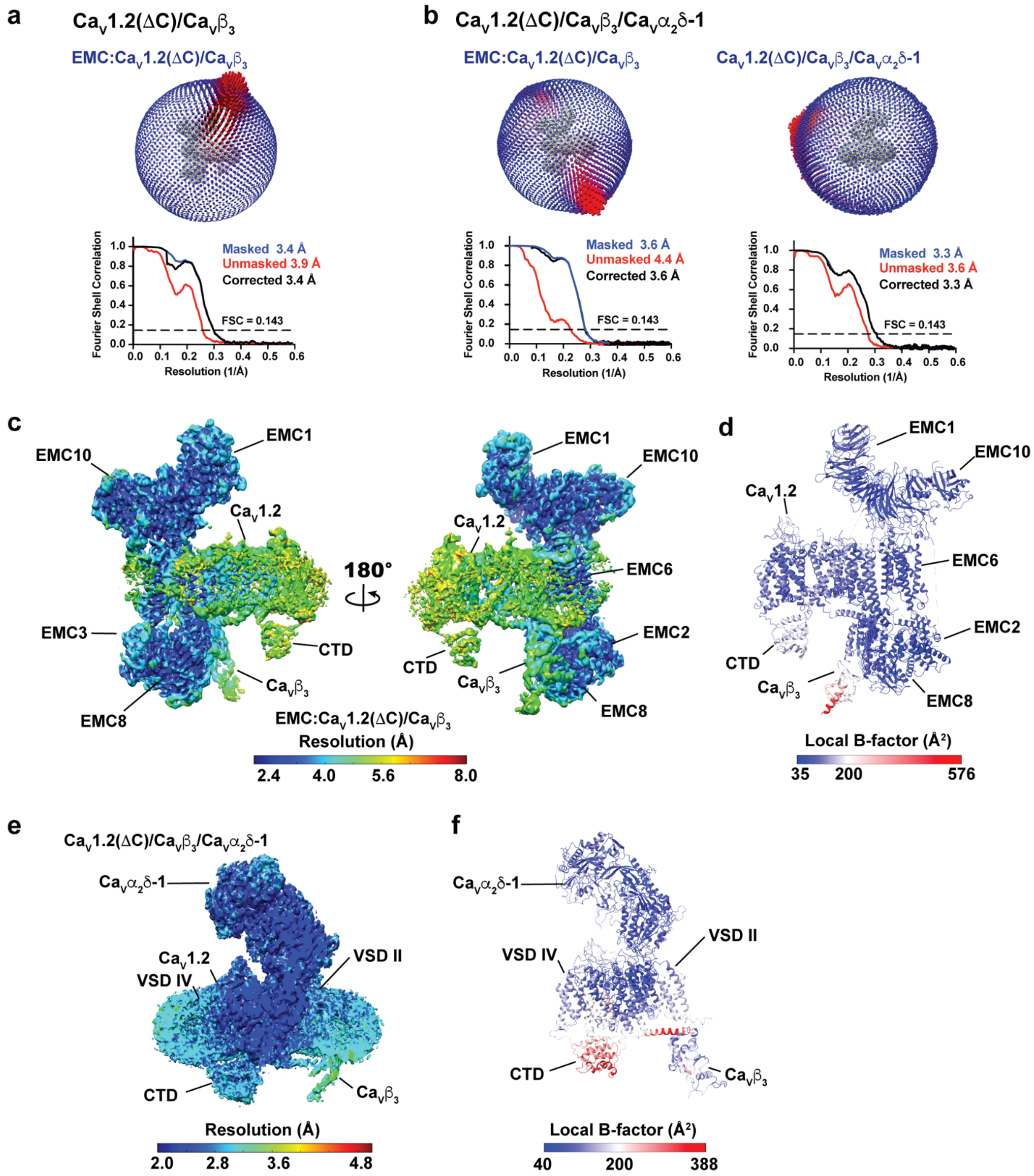


**Figure S3 EMC:CaV1.2(ΔC)/CaVβ3 and CaV1.2(ΔC)/CaV β3/CaVα2δ-1 map and model quality** **a-b**, Particle distribution plots and gold-standard Fourier shell correlation (FSC) curves for the EMC:CaV1.2(ΔC)/CaVβ3 complex from **a,** CaV1.2(ΔC)//CaVβ3 and **b,** CaV1.2(ΔC)/CaVβ3/CaVα2δ-1 samples and **b,** CaV1.2(ΔC)/CaVβ3/CaVα2δ-1 complex from the CaV1.2(ΔC)/CaVβ3/CaVα2δ-1 sample. **c-d**, EMC:$Ca_V$1.2(ΔC)/$Ca_V\beta_3$ complex **c,** local resolution and **d,** local B-factor. **e-f**, $Ca_V$1.2(ΔC)/$Ca_V\beta_3$/$Ca_V\alpha_2\delta$-1 complex **e,** local resolution and **f,** local B-factor. Select elements of each complex are labeled.

**Figure S4** **Chen *et al.***

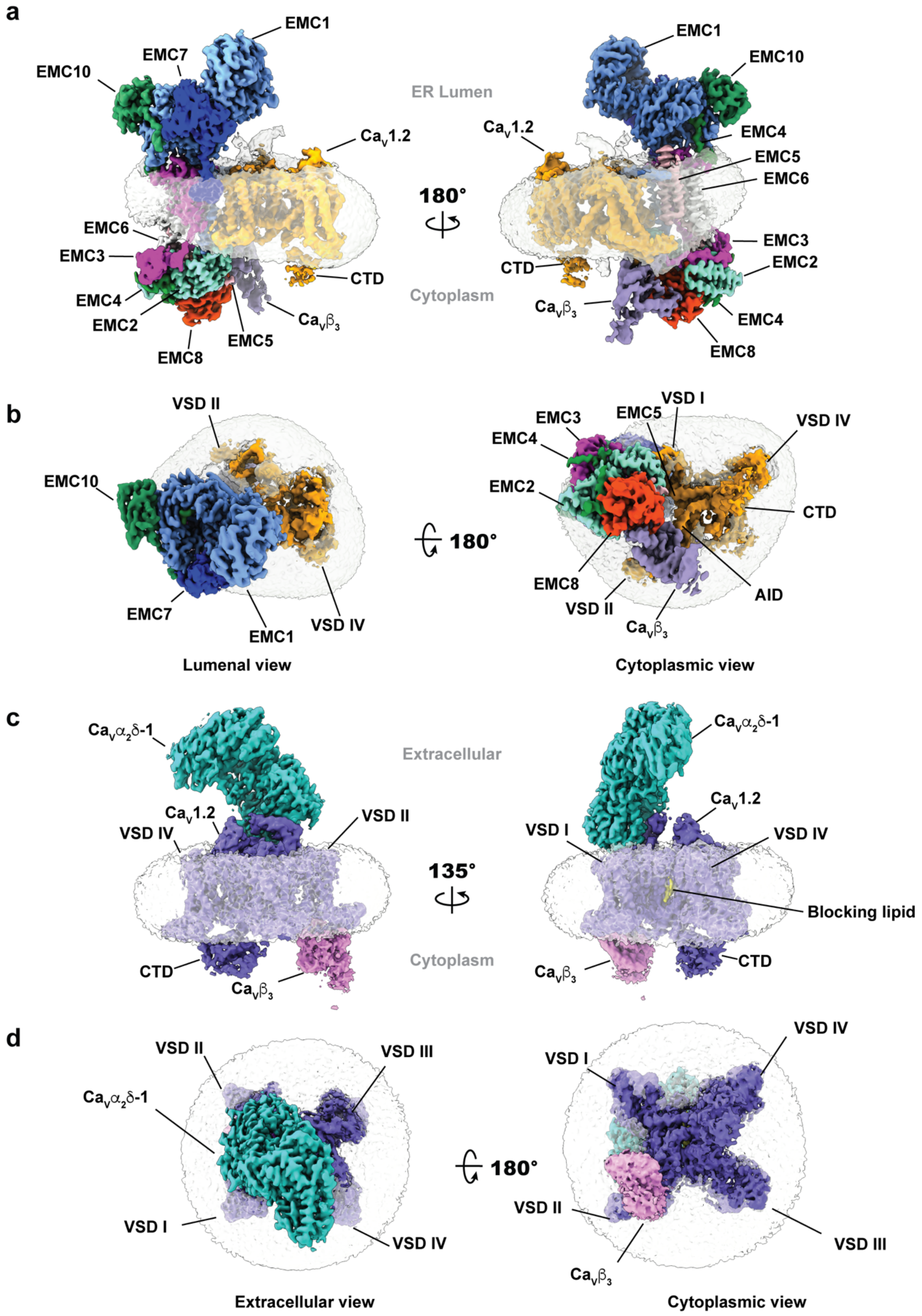

**Figure S4 Cryo-EM maps of EMC:$Ca_V$1.2(ΔC)/$Ca_V\beta_3$ and $Ca_V$1.2(ΔC)/$Ca_V\beta_3$/$Ca_V\alpha_2\delta$-1 complexes.** EMC:$Ca_V$1.2(ΔC)/$Ca_V\beta_3$ complex **a,** side views and **b,** lumenal (left) and cytoplasmic (right) views. Subunits are colored as: EMC1 (light blue), EMC2 (aquamarine), EMC3 (light magenta), EMC4 (Forest), EMC5 (light pink), EMC6 (white), EMC7 (marine), EMC8 (orange), EMC10 (smudge), $Ca_V$1.2 (bright orange), and $Ca_V\beta_3$ (lavender). $Ca_V$1.2(ΔC)/$Ca_V\beta_3$/$Ca_V\alpha_2\delta$-1 **c,** side views and **d,** extracellular (left) and cytoplasmic (right) views. Subunits are colored as: $Ca_V$1.2 (slate), $Ca_V\beta_3$ (violet), and $Ca_V\alpha_2\delta$-1 (greencyan). Detergent micelle is clear.

### Figure S5

12 sep Chen *et al.*

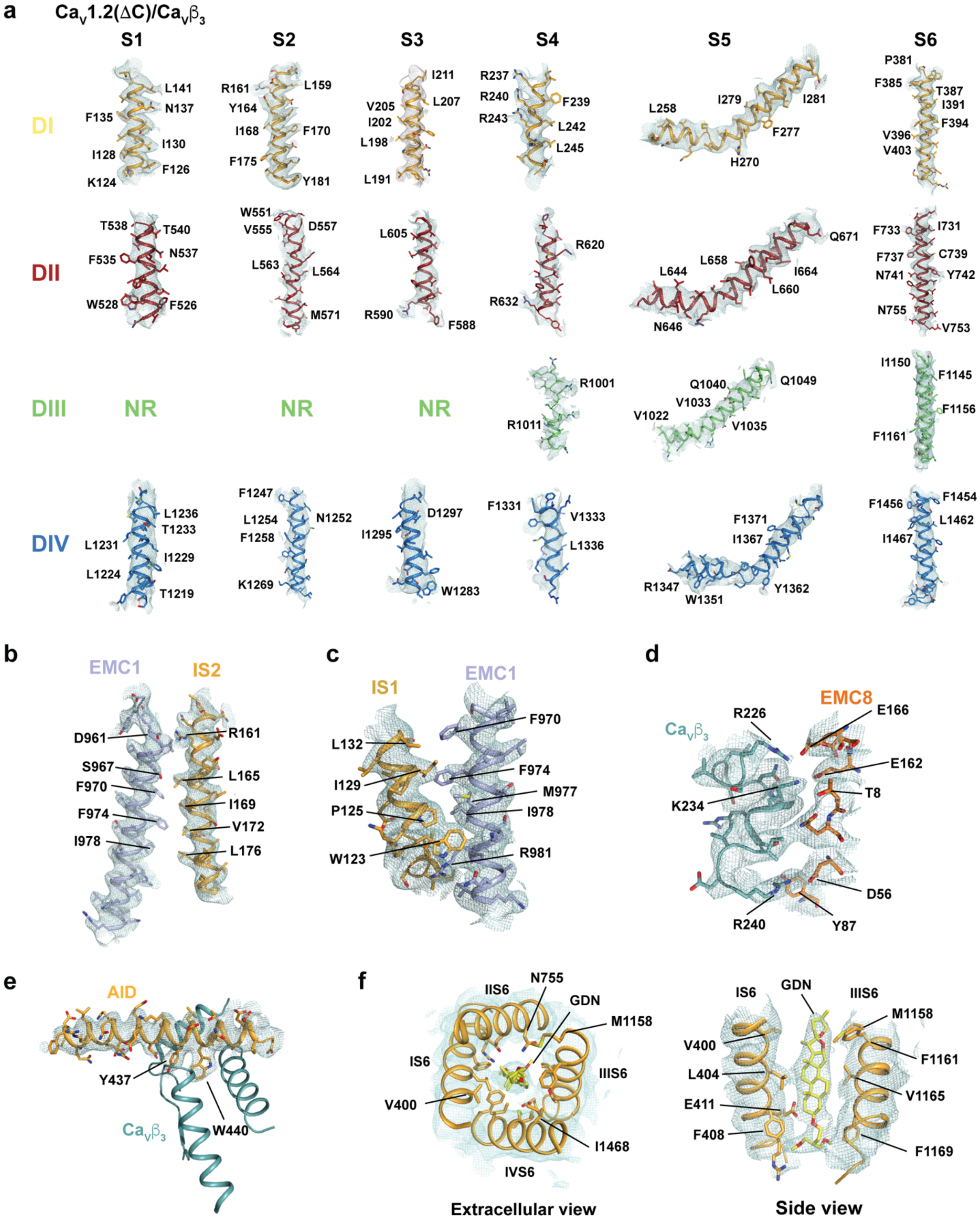


**Figure S5 Cryo-EM maps for EMC:CaV1.2(ΔC)/CaVβ3 complex elements**. **a,** Electron microscopy maps for the indicated elements from CaV1.2. Domains are colored I (yellow orange),

II (firebrick), III (lime), and IV (marine) ‘NR’ indicates not resolved. Maps are rendered at 4-5$\sigma$ **.** **b,** TM dock -IS2 **c,** TM dock -IS1 **d**, Cyto dock, **e,** AID helix, and **f,** $Ca_V$1.2 pore-blocking GDN and surrounding S6 helices.

**Figure S6** **Chen *et al.***

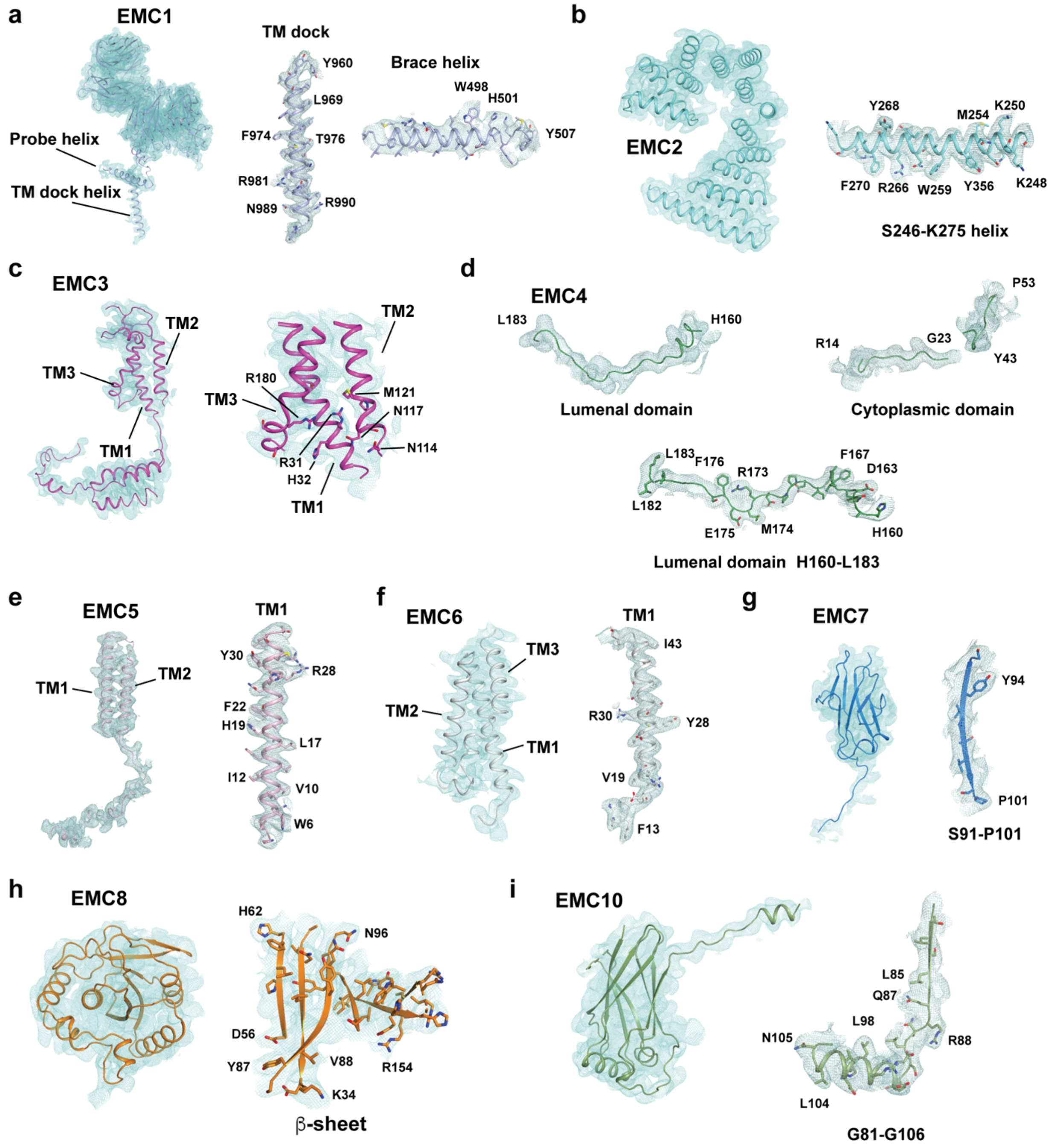


**Figure S6 Cryo-EM maps for representative EMC elements of the EMC:Ca$_V$1.2(ΔC)/Ca$_V$β$_3$ complex**. **a-i,** Cryo-EM maps for the indicated EMC subunits from the EMC:Ca$_V$1.2(ΔC)/Ca$_V$β$_3$ complex and select segments. Maps are rendered at 4-5σ**.**

**Figure S7** **Chen *et al.***

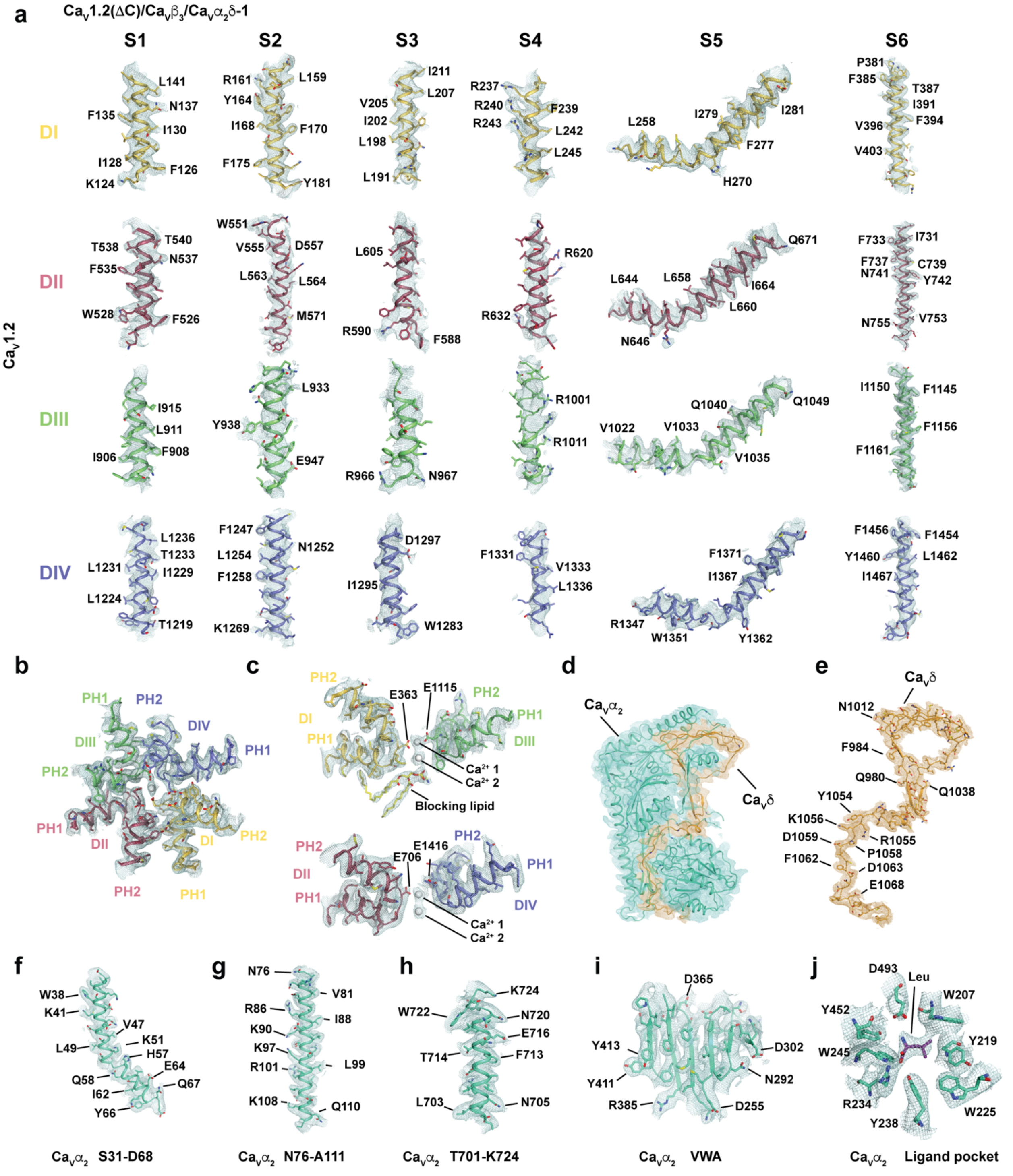


**Figure S7 Cryo-EM maps for representative segments of the Ca$_V$1.2(ΔC)/Ca$_V$β$_3$/Ca$_V$α$_2$δ-1 complex**. **a,** Cryo-EM maps for the indicated elements from Ca$_V$1.2. **b** and **c,** Electron microscopy maps for the Ca$_V$1.2 pore showing: **b,** extracellular and **c,** lateral views. Domains are colored I (yellow orange), II (raspberry), III (lime), and IV (slate). Blocking lipid (yellow) and calcium ions (white spheres) are indicated. **d-j,** Electron density for: **d,** $Ca_V\alpha_2\delta$-1 (aquamarine and sand) **e,** $Ca_V\delta$, **f-i,** Representative $Ca_V\alpha_2$ elements, and **j,** Leucine/gabapentin binding site. Maps are rendered at 4-5$\sigma$**.**

**Figure S8** **Chen *et al.***

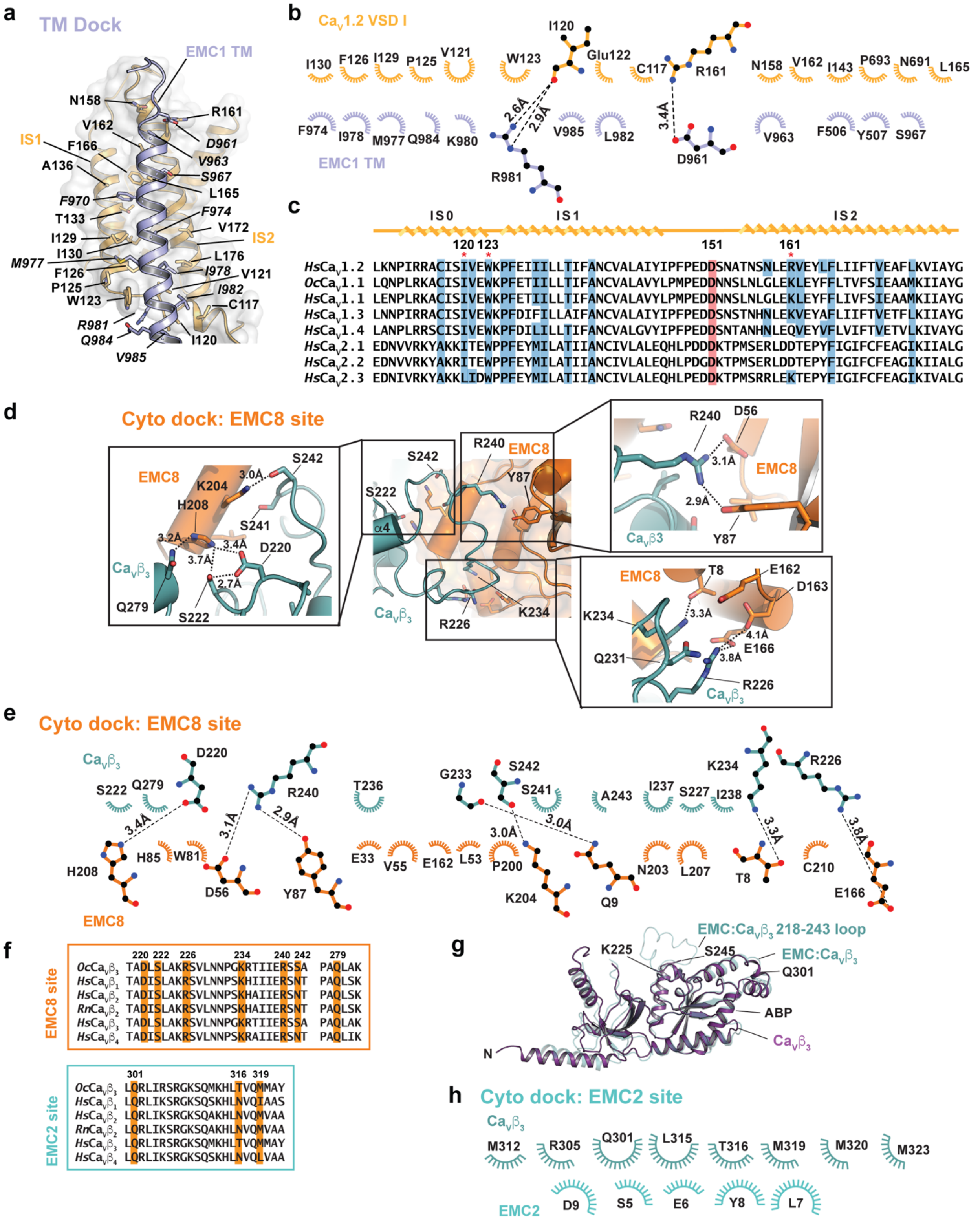


**Figure S8 EMC:CaV1.2/CaVβ3 binding sites**. **a,** View of the TM dock interaction. EMC1 TM1 (slate) and CaV1.2 VSD I (yellow orange) interface. Interface buries 1051 Å$^2$. Select elements and residues are indicated. EMC1 residues are in italics. **b,** LigPLOT (*1*) diagram of EMC1 TM:$Ca_V$1.2 VSD I interactions showing ionic interactions (dashed lines) and van der Waals contacts ≤ 5Å. **c,** Sequence comparison of the indicated VSDI sequences for human $Ca_V$1.2 (*Hs*$Ca_V$1.2 (109-182)) (Uniprot Q13936-20) with rabbit $Ca_V$1.1 (*Oc*$Ca_V$1.1 (36-109)) (NCBI: NP_001095190.1), and human L-type (*Hs*$Ca_V$1.1 (36-109), *Hs*$Ca_V$1.3 (111-184), and *Hs*$Ca_V$1.4 (77-150)) (NCBI: NP_000060.2, NP_000711.1, and NP_005174.2) and non-L-Type (*Hs*$Ca_V$2.1 (83-156), *Hs*$Ca_V$2.2 (80-153), and *Hs*$Ca_V$2.3 (74-147)) (NCBI: NP_000059.3, NP_000709.1, and NP_001192222.1) channels. Red asterisks indicate residues involved in the cation-$\pi$ pocket (120 and 123) and salt bridge (161). Red band highlights the residue that coordinates the $Ca^{2+}$ ion in the $Ca_V\alpha_2\delta$ VWA domain. **d,** $Ca_V\beta_3$:EMC8 interaction. Callouts show the details of the indicated parts of the $Ca_V\beta_3$ NK loop interaction with EMC8. **e,** LigPLOT (*1*) diagram of $Ca_V\beta_3$:EMC8 interactions showing ionic interactions (dashed lines) and van der Waals contacts ≤ 5Å. **f**, Sequence conservation for the indicated $Ca_V\beta$ elements from the EMC8 (top) and EMC2 (bottom) interaction sites**.** *Oc*$Ca_V\beta_3$ (Uniprot P54286; 218-243, 277-282; 300-322); *Hs*$Ca_V\beta_1$ (Uniprot Q02641.3; 270-295, 329-334; 352-374); *Hs*$Ca_V\beta_2$ (Uniprot Q08289; 322-347, 381-386; 404-426); *Rn*$Ca_V\beta_2$ (Uniprot Q8VGC3; 318-343, 377-382; 400-422); *Hs*$Ca_V\beta_3$ (Uniprot P54284; 218-243, 277-282; 300-322); *Hs*$Ca_V\beta_4$ (Uniprot O00305; 260-285, 319-324; 342-364). **g,** Superposition of rat $Ca_V\beta_3$ alone (violet, PDB:1VYU, chain B (*2*)) and $Ca_V\beta_3$ from the EMC complex. Boundaries of the disordered part of the T218-A243 loop in $Ca_V\beta_3$, and Q301, and ABP, are indicated, ($RMSD_{C\alpha}$ = 1.39Å). **h,** LigPLOT (*1*) diagram of $Ca_V\beta_3$:EMC2 interactions showing van der Waals contacts ≤ 5Å.

### Figure S9 

**Chen *et al.***

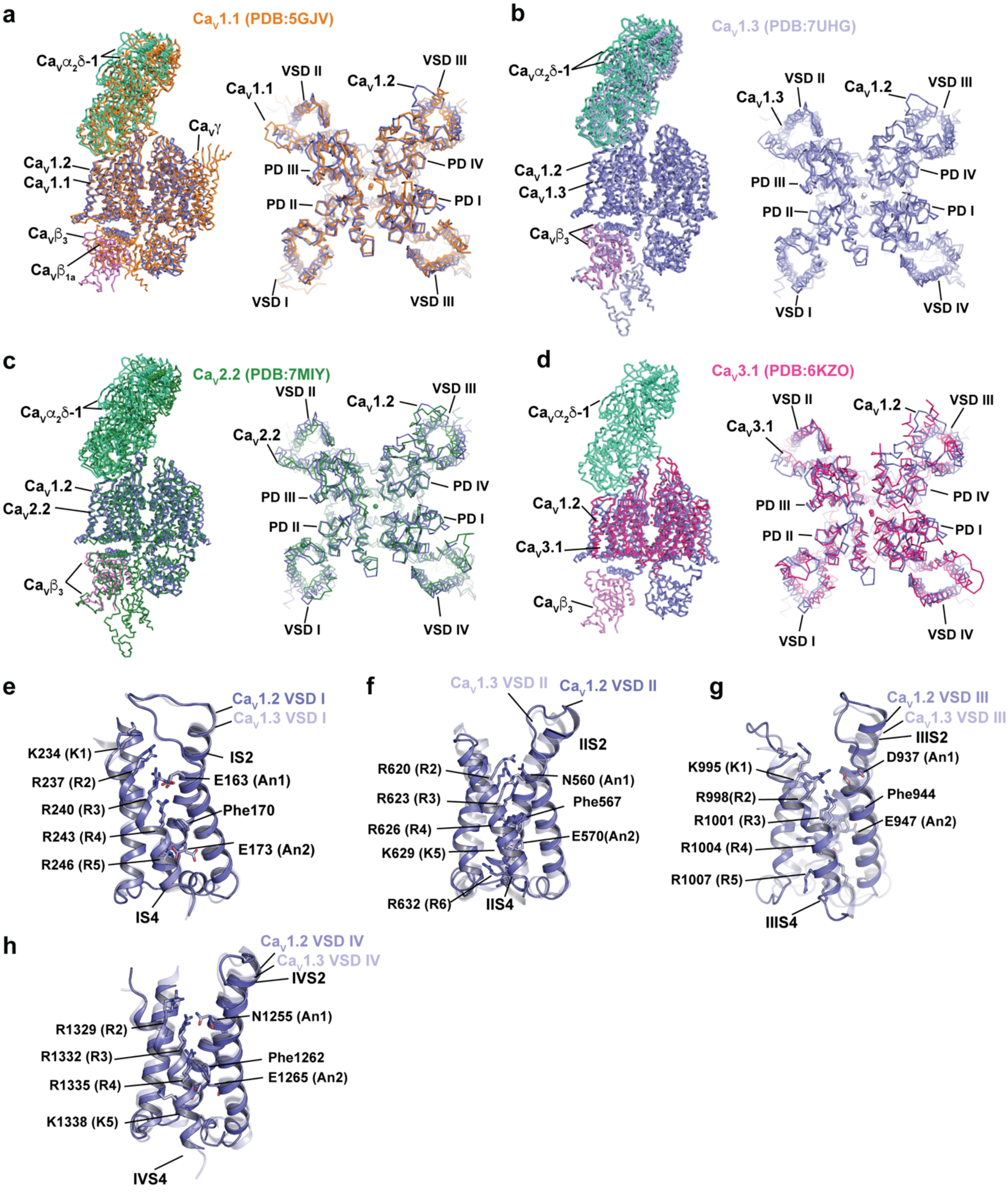


**Figure S9 Structural comparison of the $Ca_V1.2(\Delta C)/Ca_V\beta_3/Ca_V\alpha_2\delta$-1 complex with other $Ca_V$s. a-d,** Superposition of $Ca_V1.2(\Delta C)/Ca_V\beta_3/Ca_V\alpha_2\delta$-1 with **a,** rabbit $Ca_V1.1$ (PDB:5GJV)(*3*) (orange), **b,** human $Ca_V1.3$ (PDB:7UHG) (*4*) (light blue), **c,** human $Ca_V2.2$ (PDB:7MIY)(*5*) (forest), and **d,** human $Ca_V3.1$ (PDB:6KZO) (*6*)(hot pink). $Ca_V1.2$ complex subunits are colored: $Ca_V1.2$ (slate), $Ca_V\beta_3$ (violet), and $Ca_V\alpha2\delta$ (greencyan). Channel elements are indicated. **e-h,** Comparison of the indicated $Ca_V1.2$ VSDs from the $Ca_V1.2(\Delta C)/Ca_V\beta3/Ca_V\alpha_2\delta$-1complex (slate) with the corresponding VSDs (light blue) $Ca_V1.3$ (PDB:7UHG) (*4*). Gating charge residues, anionic counter charges (An1 and An2) and aromatic site of the charge transfer center (*3, 5, 7*) are shown. $RMSD_{C\alpha}$ = 0.67 Å, 2.29 Å, 4.19 Å, and 2.45 Å for VSD I, VSD II, VSD III, and VSD IV, respectively.

**Figure S10** **Chen *et al.***

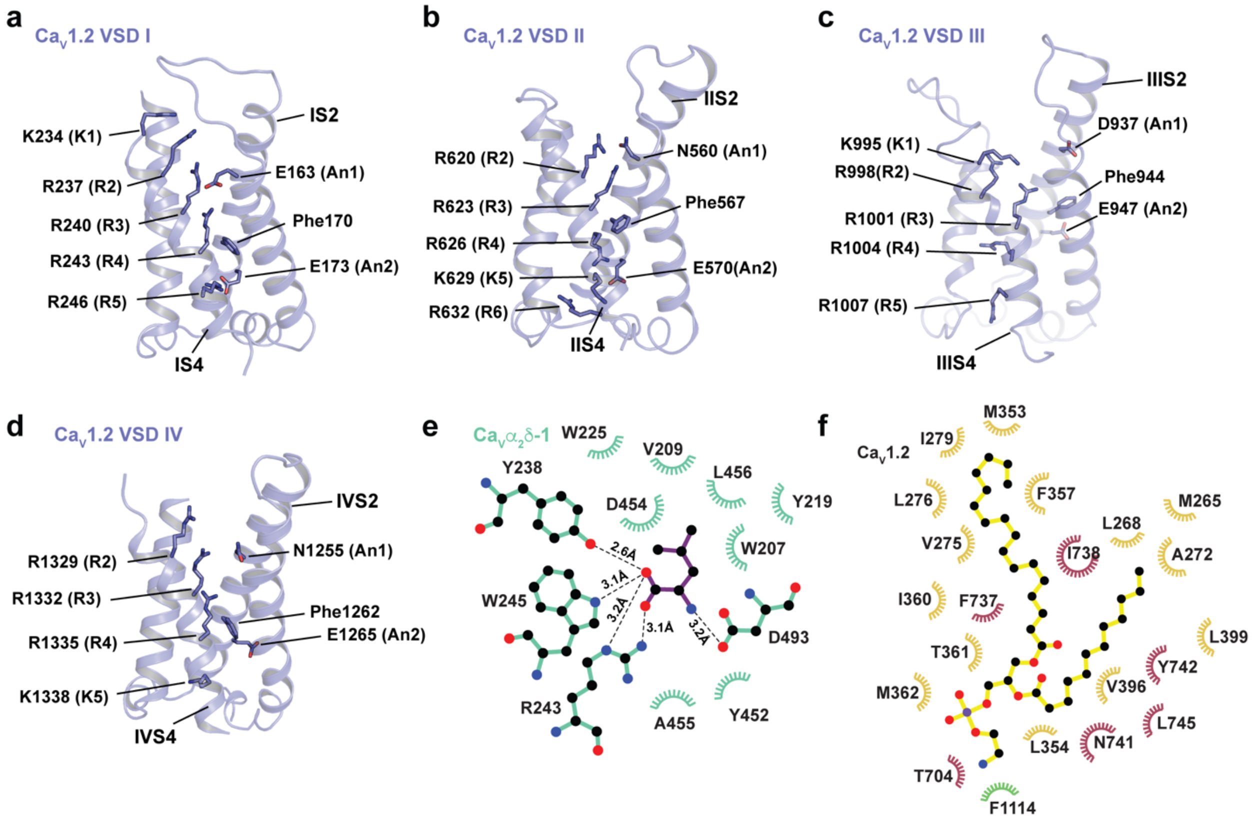


**Figure S10 CaV1.2 structural details. a-d,** Structures of the indicated VSDs from the CaV1.2(ΔC)/CaVβ3/CaVα2δ-1 complex. Gating charge residues, anionic counter charges (An1 and An2) and aromatic site of the charge transfer center (*3, 5, 7*) are shown. **e,** LigPLOT (*1*) diagram of the CaVα2δ-1 leucine binding site showing hydrogen bonds and ionic interactions (dashed lines) and van der Waals contacts ≤ 5Å. **f,** LigPLOT (*1*) diagram of blocking lipid: CaV1.2 showing van der Waals contacts ≤ 5Å. Domain I (yellow orange), Domain II (dark red), and Domain III (green) residues are indicated.

**Figure S11** **Chen *et al.***

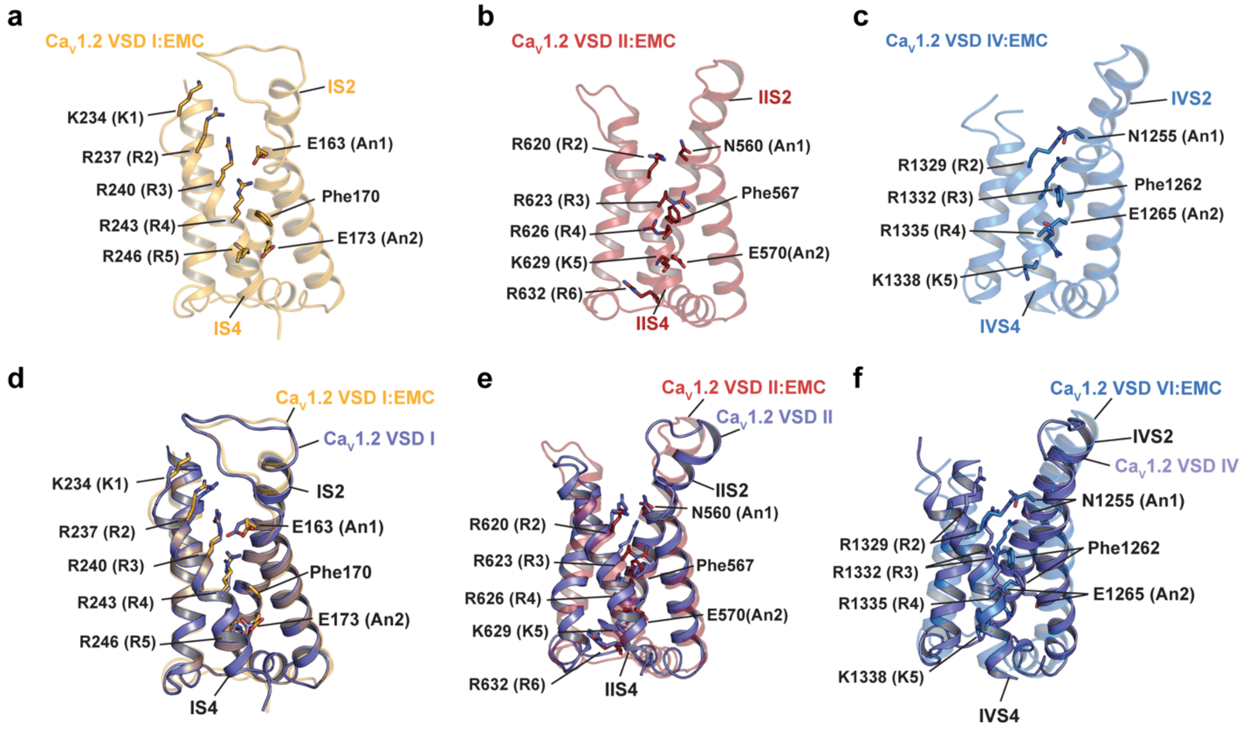


**Figure S11 VSD domains from the EMC:CaV1.2/CaVβ3 complex**. Structures of **a,** VSD I, **b,** VSD II, and **c,** VSD IV from the EMC1:CaV1.2/CaVβ3 complex and **d-f,** comparison with the corresponding VSDs (slate) from the CaV1.2/CaVβ3/CaVα2δ-1 complex. Gating charge residues, anionic counter charges (An1 and An2) and aromatic site of the charge transfer center (*3, 5, 7*) are shown.

**Figure S12** **Chen *et al.***

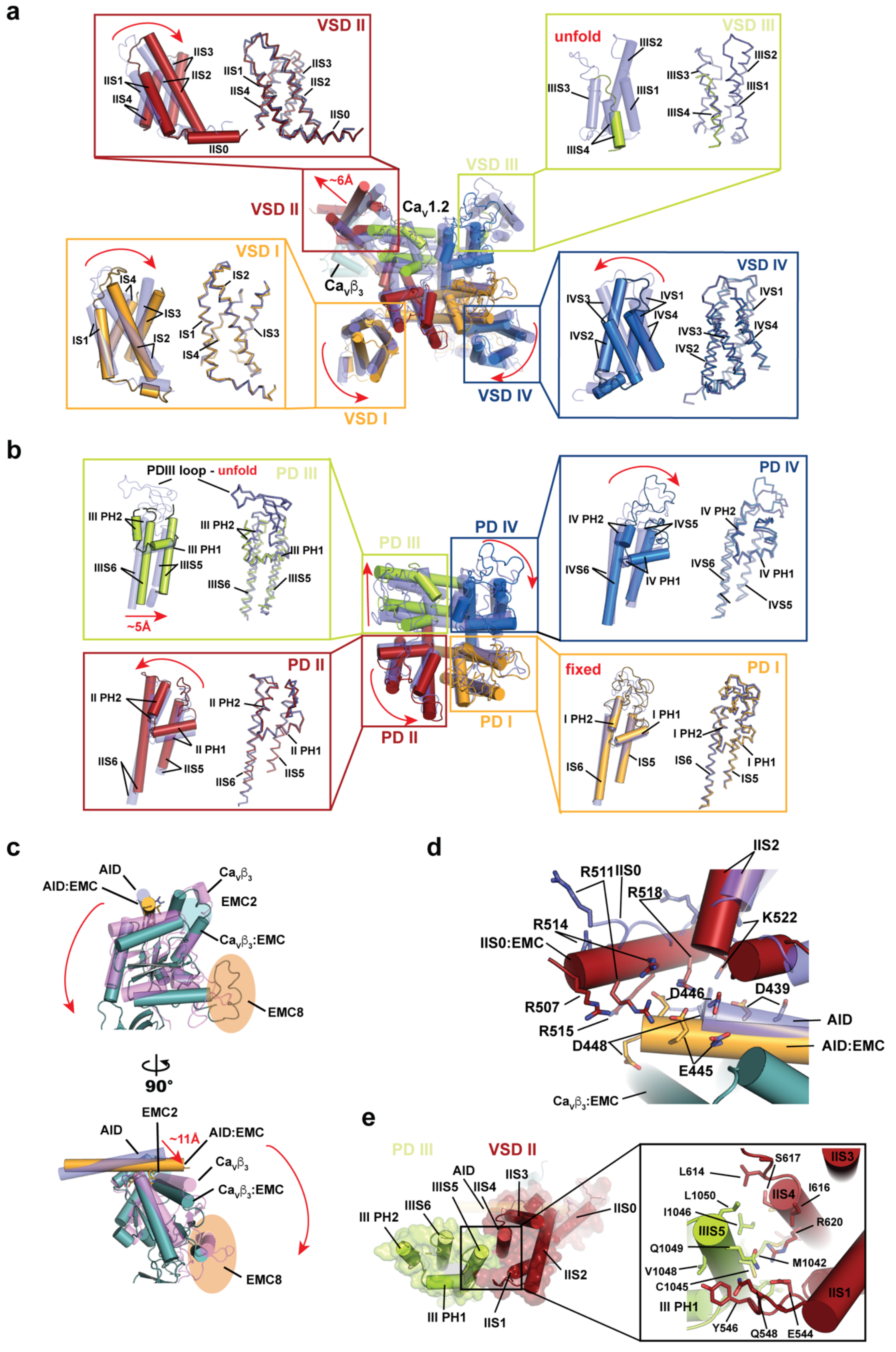

**Figure S12 Conformational changes between EMC-bound and $Ca_V\alpha_2\delta$-bound $Ca_V1.2/Ca_V\beta_3$.** Superposition of $Ca_V1.2$ from the EMC:$Ca_V1.2(\Delta C)/Ca_V\beta_3$ and $Ca_V1.2(\Delta C)/Ca_V\beta_3/Ca_V\alpha_2\delta$-1 complexes showing **a,** VSD conformational changes. **b,** PD conformational changes showing the superposition from 'a'. Insets show each PD. Elements $Ca_V1.2$ from the EMC complex are: VSD I/PD I (yellow orange), VSD II/PD II (firebrick), VSD III/PD III (lime), VSD IV/PD IV (marine). $Ca_V1.2$ (slate) and $Ca_V\beta_3$ (violet) from $Ca_V1.2/Ca_V\beta_3/Ca_V\alpha_2\delta$-1 and $Ca_V\beta_3$ from the EMC (light teal) are semi-transparent. **c,** Superposition of $Ca_V\beta_3$ and the $Ca_V1.2$ AID helix from the EMC:$Ca_V1.2(\Delta C)/Ca_V\beta_3$ and $Ca_V1.2(\Delta C)/Ca_V\beta_3/Ca_V\alpha_2\delta$-1 complexes. Location of EMC8 is indicated by the orange oval. Red arrows in 'a-c' indicate conformational changes between $Ca_V1.2/Ca_V\beta_3/Ca_V\alpha_2\delta$ and EMC:$Ca_V1.2/Ca_V\beta_3$. **d,** Comparison of IIS0 and surrounding regions in the EMC complex (VSDII, firebrick; AID (yellow orange), and $Ca_V\beta_3$ (light teal)) their corresponding elements in $Ca_V1.2/Ca_V\beta_3/Ca_V\alpha_2\delta$-1 (slate). **e,** Interactions between VSD II:PD III in the $Ca_V\beta$:AID:VSD II:PD III subcomplex from the EMC:$Ca_V1.2(\Delta C)/Ca_V\beta_3$ structure.

**Figure S13** **Chen *et al.***

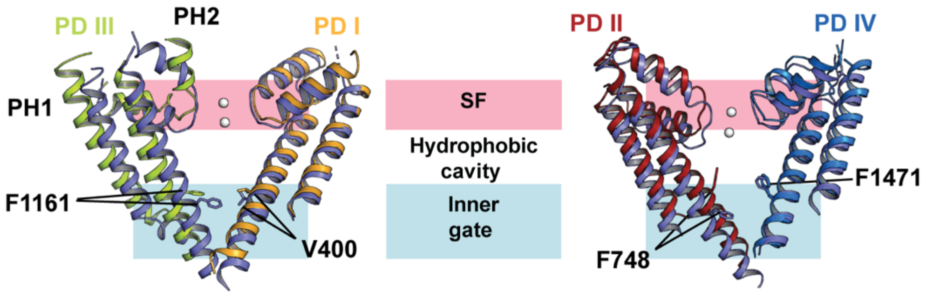


**Figure S13 $Ca_V1.2$ pore domain superposition for EMC-bound and $Ca_V\alpha_2\delta$-bound $Ca_V1.2/Ca_V\beta_3$.** Pore domains from the EMC-bound complex are: PD I (yellow orange), PD II (firebrick), PD III (lime), and PD IV (marine). Pore domains from $Ca_V1.2(\Delta C)/Ca_V\beta_3/Ca_V\alpha_2\delta$-1 are slate. Calcium ions are from $Ca_V1.2(\Delta C)/Ca_V\beta_3/Ca_V\alpha_2\delta$-1. Selectivity filter (SF), hydrophobic cavity, and inner gate regions and select residues are indicated.

**Figure S14** **Chen *et al.***

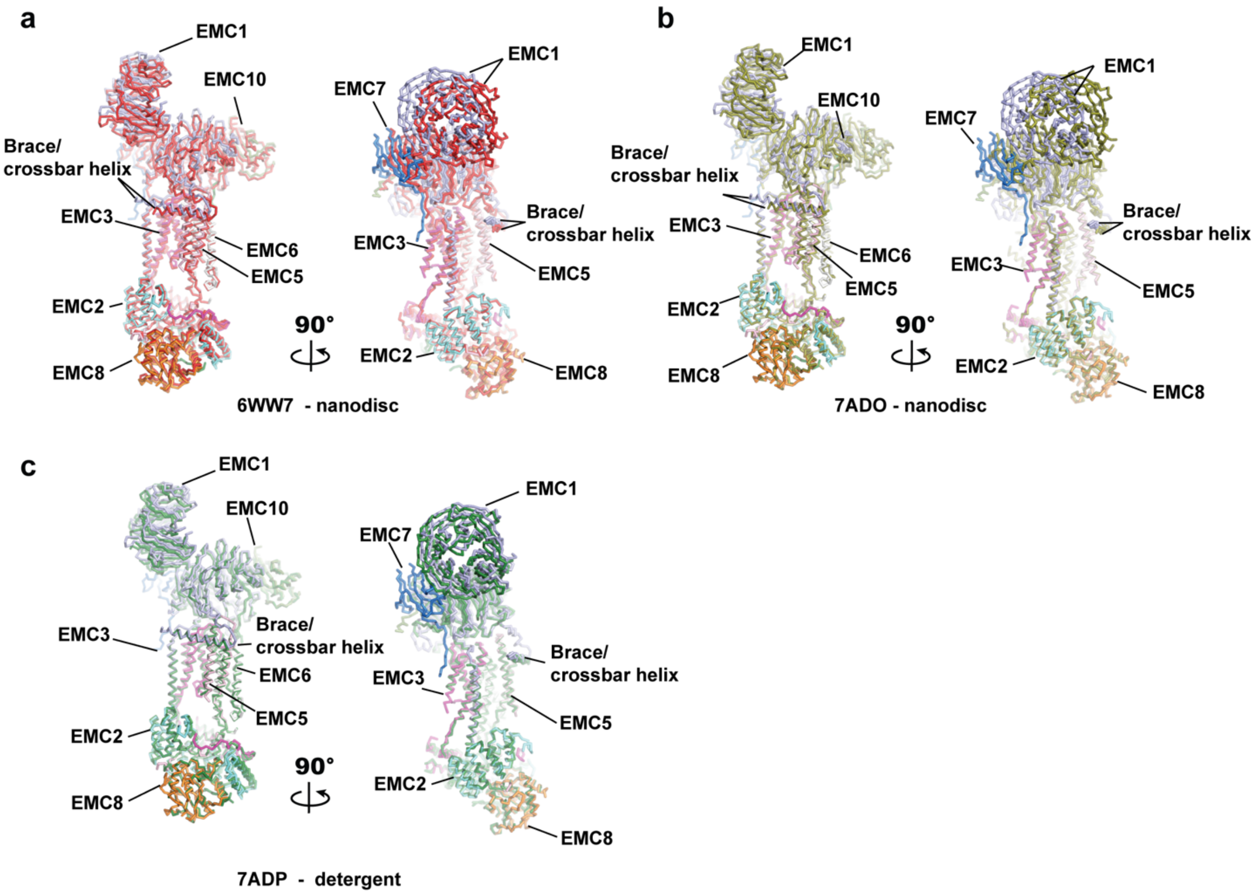


**Figure S14 Comparison of client-loaded EMC and apo-EMC complexe*s.*** Comparison of the EMC complex from the EMC:$Ca_V1.2(\Delta C)/Ca_V\beta_3$ complex with human EMC structures **a,** EMC in lipid nanodiscs (PDB:6WW7) (*8*) (red), **b,** EMC in lipid nanodiscs (PDB:7ADO) (deep olive) (*9*), and **c,** EMC in detergent (PDB:7ADP) (forest) (*9*).

### Figure S15 

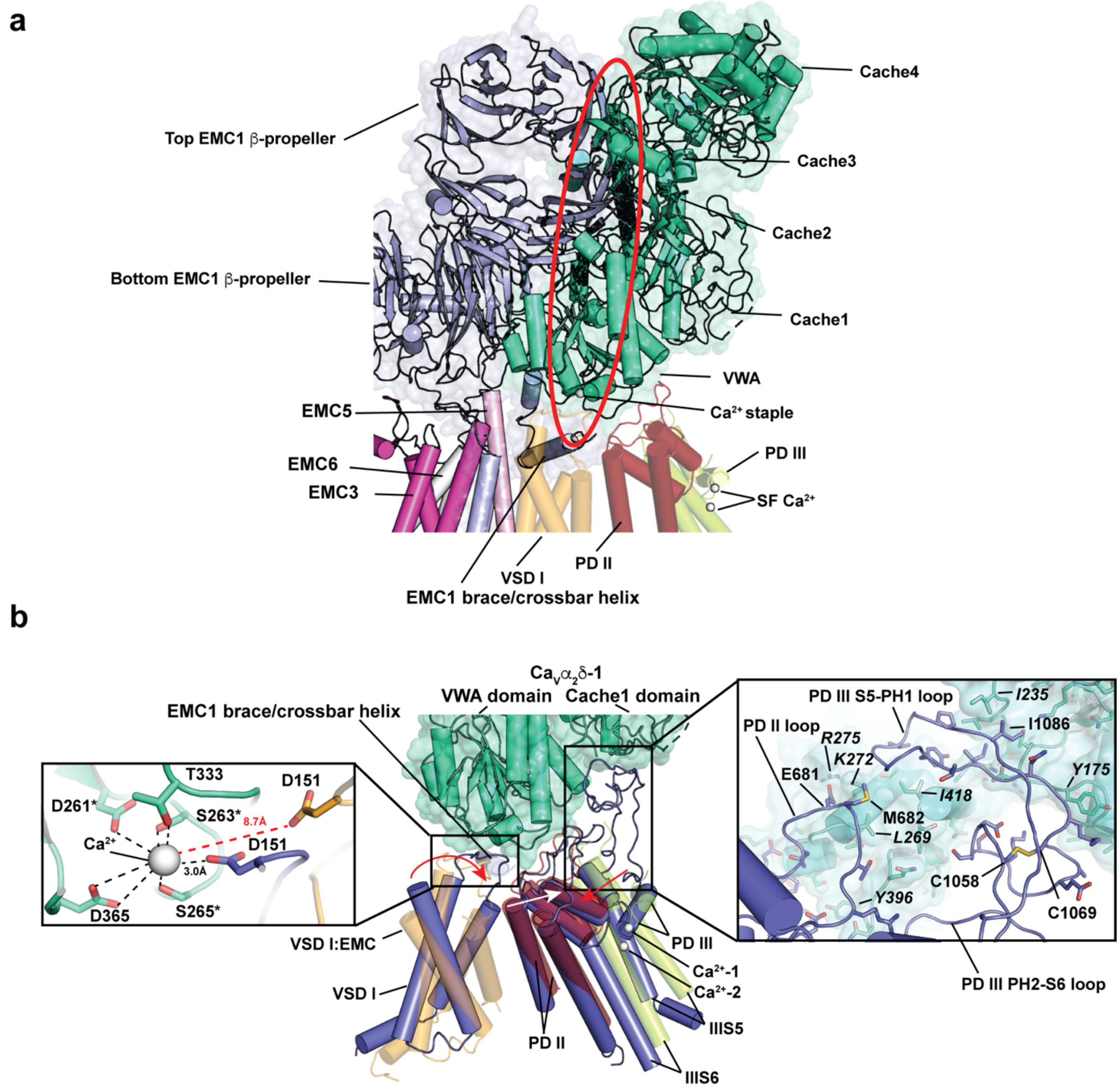


**Figure S15 EMC1/ $Ca_V\alpha_2\delta$ clash and ordering of the $Ca_V$1.2 pore and VSDs by $Ca_V\alpha_2\delta$.** **a,** Close up view of clash between EMC1 (light blue) and $Ca_V\alpha_2\delta$-1 (aquamarine). EMC1, EMC3 (magenta), EMC5 (pink), EMC6 (white), VSD I (yellow orange), PD II (firebrick), and PD III (lime) from the EMC:$Ca_V$1.2/$Ca_V\beta_3$ complex are shown. Red oval highlights clash regions. $Ca_V\alpha_2\delta$-1 domains are indicated. $Ca^{2+}$ staple is indicated. SF calcium ions are from $Ca_V$1.2(ΔC)/$Ca_V\beta_3$/$Ca_V\alpha_2\delta$-1 and mark the location of the pore in the $Ca_V\alpha_2\delta$-assembled channel. **b,** Superposition of VSD I (yellow orange), PD II (firebrick), and PD III (lime) from the EMC complex (semi-transparent) and their corresponding parts from $Ca_V$1.2(ΔC)/$Ca_V\beta_3$/$Ca_V\alpha_2\delta$-1 (slate). $Ca_V\alpha_2\delta$-1 (aquamarine) is shown as a semi-transparent surface. Calcium ions are from $Ca_V$1.2(ΔC)/$Ca_V\beta_3$/$Ca_V\alpha_2\delta$-1. Left inset shows the coordination of the $Ca^{2+}$ staple by the VWA domain MIDAS and D151 in $Ca_V$1.2(ΔC)/$Ca_V\beta_3$/$Ca_V\alpha_2\delta$-1. Red distance shows the position of $Ca_V$1.2 D151 in the EMC complex relative to the calcium ion in the $Ca_V\alpha_2\delta$ complex. Asterisks mark positions where coordinated alanine mutation impair the ability of $Ca_V\alpha_2\delta$ to enhance $Ca_V$ currents and surface expression (*10*). Right inset shows the extensive contacts between PD II and PD III loops with $Ca_V\alpha_2\delta$ in the $Ca_V$1.2(ΔC)/$Ca_V\beta_3$/$Ca_V\alpha_2\delta$-1 complex. The PD III loops are disordered in the EMC:$Ca_V$1.2/$Ca_V\beta_3$ complex. $Ca_V\alpha_2\delta$-1 residues are in italics.

### Figure S16

**Chen *et al.***

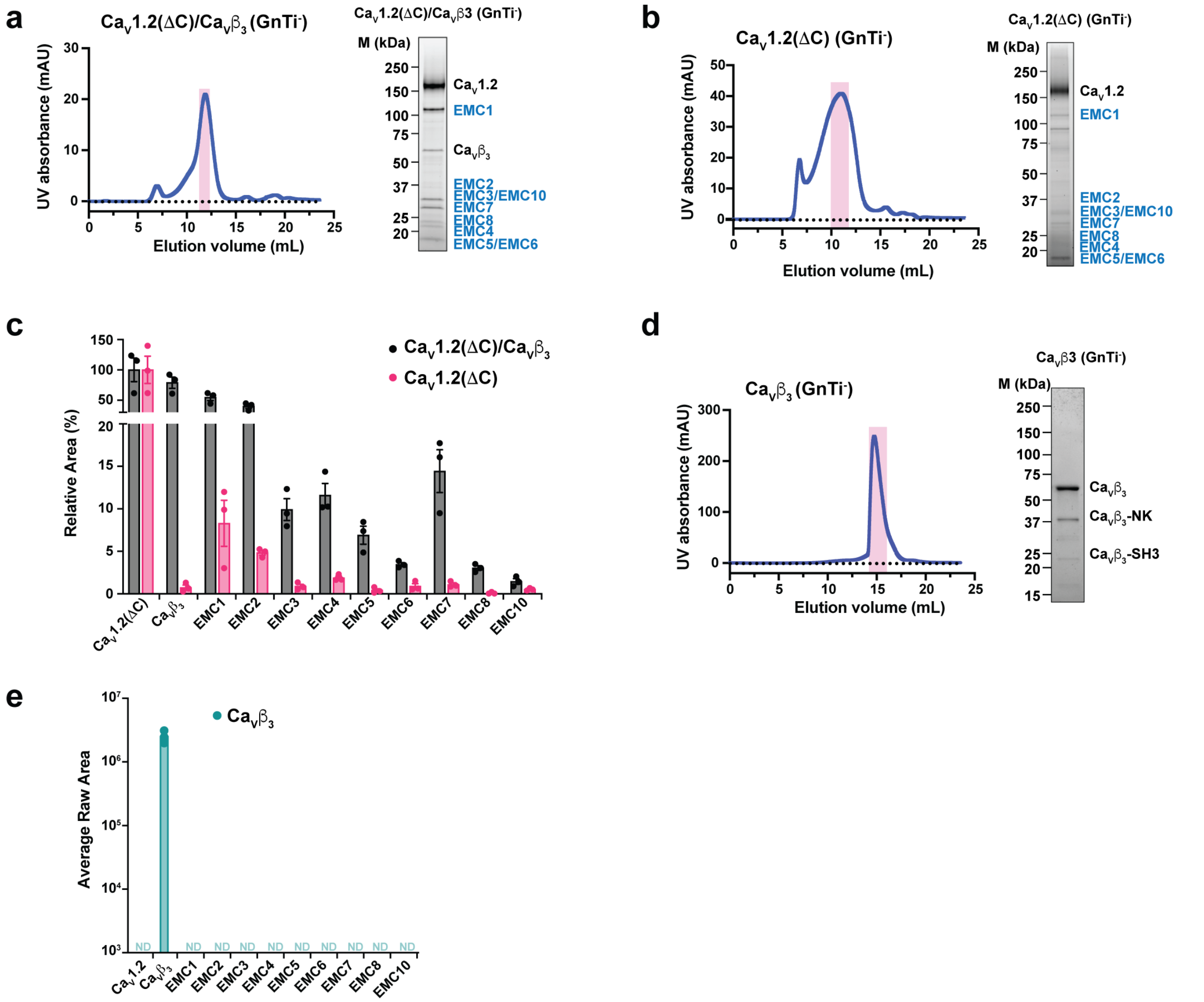


**Figure S16 Exemplar purification of $Ca_V$ subunit combinations a-c** Superose 6 Increase 10/300 GL chromatogram and peak fraction SDS-PAGE for: **a,** $Ca_V1.2(\Delta C)/Ca_V\beta_3$ **b,** $Ca_V1.2(\Delta C)$, and **c,** Relative detection by mass spectrometry of EMC proteins with respect to $Ca_V1.2(\Delta C)$ across 3 replicates. Error bars are calculated as SEM. **d,** $Ca_V\beta_3$. $Ca_V\beta_3$-NK and $Ca_V\beta_3$-SH3 are $Ca_V\beta_3$ proteolytic fragments. **e,** Absolute detection of EMC proteins and $Ca_V\beta_3$ by mass spectrometry following expression and purification of $Ca_V\beta_3$. Error bars are calculated as SEM. ND denotes not detected.

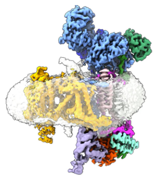

**Movie S1 Overview of the EMC:Ca$_V$1.2/Ca$_V\beta_3$ complex.** Movie shows the EMC:Ca$_V$1.2($\Delta$C)/Ca$_V\beta_3$ complex and highlights the TM and Cyto dock sties. Subunits are colored as: EMC1 (light blue), EMC2 (aquamarine), EMC3 (light magenta), EMC4 (Forest), EMC5 (light pink), EMC6 (white), EMC7 (marine), EMC8 (orange), EMC10 (smudge), Ca$_V$1.2 (bright orange), and Ca$_V\beta_3$ (lavender). Detergent micelle is clear.

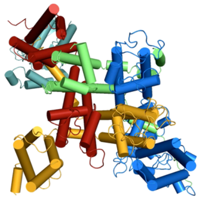

**Movie S2 Lumenal view of Ca$_V$1.2 conformational changes upon EMC binding.** Morph between Ca$_V$1.2 conformations in Ca$_V$1.2($\Delta$C)/Ca$_V\beta_3$/Ca$_V\alpha_2\delta$-1 (start and end) and the EMC:Ca$_V$1.2($\Delta$C)/Ca$_V\beta_3$ complex (middle). Ca$_V$1.2 elements are : VSD I/PD I (yellow orange), VSD II/PD II (firebrick), VSD III/PD III (lime),and VSD IV/PD IV (marine). Ca$_V\beta_3$ (light teal).

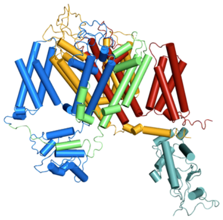

**Movie S3 Side view of Ca$_V$1.2 conformational changes upon EMC binding.** Morph between Ca$_V$1.2 conformations in the Ca$_V$1.2($\Delta$C)/Ca$_V\beta_3$/Ca$_V\alpha_2\delta$-1 (start and end) and EMC:Ca$_V$1.2($\Delta$C)/Ca$_V\beta_3$ complex (middle). Ca$_V$1.2 elements are :VSD I/PD I (yellow orange), VSD II/PD II (firebrick), VSD III/PD III (lime),and VSD IV/PD IV (marine). Ca$_V\beta_3$ (light teal). View is of the side of the VSD III/PD III S4/S5 linker.

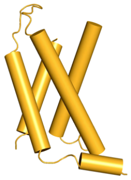

**Movie S4 $Ca_V$1.2 VSD I conformational changes upon EMC binding.** Morph of VSD I conformations in the $Ca_V1.2(\Delta C)/Ca_V\beta_3/Ca_V\alpha_2\delta$-1 (start and end) and EMC:$Ca_V1.2(\Delta C)/Ca_V\beta_3$ complexes (middle). View is of the IS1/IS2 groove that binds the EMC 1 TM dock (cf. Fig. S8a and S12a).

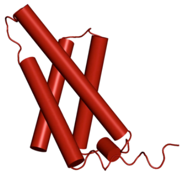

**Movie S5 $Ca_V$1.2 VSD II conformational changes upon EMC binding.** Morph of VSD II conformations in the $Ca_V1.2(\Delta C)/Ca_V\beta_3/Ca_V\alpha_2\delta$-1 (start and end) and EMC:$Ca_V1.2(\Delta C)/Ca_V\beta_3$ complexes (middle). View is onto IIS0, IIS1, and IIS2 (cf. Fig. S12a).

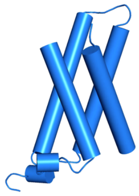

**Movie S6 $Ca_V$1.2 VSD IV conformational changes upon EMC binding.** Morph of VSD IV conformations in the $Ca_V1.2(\Delta C)/Ca_V\beta_3/Ca_V\alpha_2\delta$-1 (start and end) and EMC:$Ca_V1.2(\Delta C)/Ca_V\beta_3$ complexes (middle). View is onto IVS3 and IVS4 (cf. Fig. S12a).

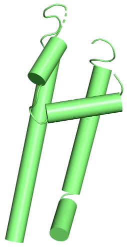

**Movie S7 $Ca_V$1.2 PD III conformational changes upon EMC binding.** Morph of PD III conformations in the $Ca_V1.2(\Delta C)/Ca_V\beta_3/Ca_V\alpha_2\delta$-1 (start and end) and EMC:$Ca_V1.2(\Delta C)/Ca_V\beta_3$ complexes (middle). View is onto the SF and pore helices (cf. Fig. S12b).

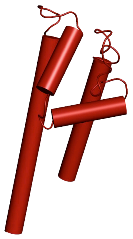

**Movie S8 $Ca_V1.2$ PD II conformational changes upon EMC binding.** Morph of PD II conformations in the $Ca_V1.2(\Delta C)/Ca_V\beta_3/Ca_V\alpha_2\delta$-1 (start and end) and EMC:$Ca_V1.2(\Delta C)/Ca_V\beta_3$ complexes (middle). View is onto the SF and pore helices (cf. Fig. S12b).

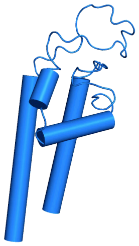

**Movie S9 $Ca_V1.2$ PD IV conformational changes upon EMC binding.** Morph of PD IV conformations in the $Ca_V1.2(\Delta C)/Ca_V\beta_3/Ca_V\alpha_2\delta$-1 (start and end) and EMC:$Ca_V1.2(\Delta C)/Ca_V\beta_3$ complexes (middle). View is onto the SF and pore helices (cf. Fig. S12b).

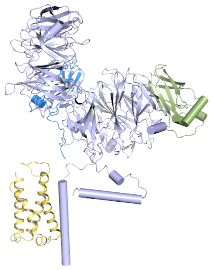

**Movie S10 EMC lumenal domain movement induced by client binding.** Morph between apo-EMC (PDB:6WW7)(*8*) (start and end) and the EMC in the EMC:$Ca_V1.2(\Delta C)/Ca_V\beta_3$ complex (middle) showing EMC1 (light blue), EMC7 (marine) and EMC10(smudge). $Ca_V1.2$ VSD I (yellow orange) from the EMC:$Ca_V1.2/Ca_V\beta_3$ complex is shown as ribbons. View is lateral to the brace/crossbar helix (cf. Fig. 5a).

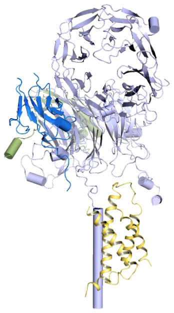

**Movie S11 EMC lumenal domain movement induced by client binding.** Morph between apo-EMC (PDB:6WW7)(*8*) (start and end) and the EMC in the EMC:$Ca_V1.2(\Delta C)/Ca_V\beta_3$ complex (middle) showing EMC1 (light blue), EMC7 (marine) and EMC10(smudge). $Ca_V1.2$ VSD I (yellow orange) from the EMC:$Ca_V1.2(\Delta C)/Ca_V\beta_3$ complex is shown as ribbons. View is axial to the brace/crossbar helix (cf. Fig. 5a).

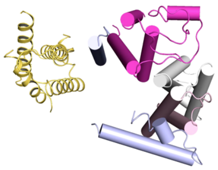

**Movie S12 EMC transmembrane domain movement induced by client binding.** Morph between apo-EMC (PDB:6WW7)(*8*) (start and end) and the EMC in the EMC in the EMC:$Ca_V1.2(\Delta C)/Ca_V\beta_3$ complex (middle) showing EMC1 (light blue), EMC3 (light magenta), EMC5 (light pink), and EMC6 (white) transmembrane helices. VSD I from the EMC:$Ca_V1.2(\Delta C)/Ca_V\beta_3$ complex is shown as ribbons (yellow orange). View is from the lumenal side (cf. Fig. 5b).

| !""#$$%&' | ()&*#%'+ ,-.# | /#'#+'-.# | 0#$")%1*%&' | 2345&6( | 7(#1*%8#$ | 79'%:;# | !)#-+ !<1=->?#*-+ @3 | !)#-+ !<1=->?#*-+ @A | !)#-+ !<1=->?#*-+ @B | C#8%-'+ !)#-+ -<1=->D#*- | !)#-+!<1=-2 E'<F+@3 | !)#-+!<1=-2 E'<F+@A | !)#-+!<1=-2 E'<F+@B | !)#-+?#*-2 E'<F+@3 | !)#-+?#*-2 E'<F+@A | !)#-+?#*-2 E'<F+@B | 7G1#"+ H5C4IA2@3 | 7G1#"+ H5C4IA2@A | 7G1#"+ H5C4IA2@B | 7G1#"+ 4JAK@3 | 7G1#"+ 4JAK@A | 7G1#"+ 4JAK@B | 7G1#"+?#*-2 E'<F2@3 | 7G1#"+?#*-2 E'<F2@A | 7G1#"+?#*-2 E'<F2@B | (LC | !M#)-6#+ C-$$ |
|---|---|---|---|---|---|---|---|---|---|---|---|---|---|---|---|---|---|---|---|---|---|---|---|---|---|---|---|
| !"#$#% | &'&"& | &'&(""& | )*+,-./01/2/31/3,450,62/47-+789:47;-33/+4<9=938,4-+2;-0"& | >?@AB#@?% | "BC | "DB | "AD$EFC% | DAB$EFC% | DA>"EFC% | DA>"EFC% | #A?DEFC% | BAD%EFC% | DA#DEFC% | | | | "@## | DD@# | D#DC | #B>C | BD%" | #>B> | | | | GH81-,8*34I. | D>@$?%A$ |
| !@(?%% | EJ&" | EJ&" | EO4:/:=N-3/42N*,/8347*:2+/H4<9=938,4" | >"#A#@?D> | $$ | $$ | $AD$EFCB | "AD$EFC% | "ADDEFC% | "ADDEFC% | "A">EFCB | #A?DEFCB | >ABDEFCB | | | | %?@ | $"% | $$D | DC# | >$@ | >@" | | | | GH81-,8*34I. | """?B$A> |
| PB>D@> | &'&Q# | &'&(Q# | )*+,-./01/2/31/3,450,62/47-+789:47;-33/+4<9=938,4=/,-0# | #$$AC?D>B | "C# | @> | "AD?EFC% | "A$#EFC% | "A?BEFC% | "A?BEFC% | "AC#EFC> | DA##EFC> | >A@$EFC> | DA>?EFC% | "A$@EFC% | #A"CEFC% | @@D | "CDD | "CDD | ""$ | "D> | "#% | @BBC | @%"C | $?"D | M/-:81-,8*34 | B>B#"A$"@ |
| (4NOP4 | QRCS | QRC | Q%.#'*%' | BATUPPANB | BT | BI | IUITS>4J | 3UOBS>4J | NUPAS>4I | 3UOBS>4J | BUTTS>4A | AUBJS>4B | IUJNS>4B | PUN3S>4B | PU43S>4B | NU3IS>4B | B34N | A4OB | 3B4A | BI | A3 | AI | 3BI | 3JI | 3OJ | 0#-.%8-*%&' | JBOJ3UOPC |
| (4OPBB | S,E! | S,E3 | !<1=-2#'&<-$# | B3NUO4BPB | AB | 3N | AUB3S>4J | BUA4S>4I | 3UI4S>4I | BUA4S>4I | TUP3S>4I | NUNOS>4B | OUTNS>4B | IU3BS>4B | 3U3JS>4B | NUOPS>4A | 33BT | IBN | A3N | TJ4 | APJ | A34 | 3NN | PI | 3AB | [-)D-.%8&.#* | IP3ONUTJP |
| (APNAI | [!5\ | [!,\ | [-<'#Z%' | B3OU3BIAN | BT | BT | AUJ4S>4I | BUIBS>4I | BUIBS>4I | BUIBS>4I | BU3TS>4I | 3U3TS>4J | 3UTOS>4J | | | BUPIS>4A | T3 | 343 | 3AT | 34B | AII | AAA | | | 3 | 0#-.%8-*%&' | OPJONU |
| (JIANT | [!A03 | [![,!A03 | Q&<*-6#28#1#'8#'*+"-<"%;.+"=-"#<+$;D;'%*+-<1=-2A]8#<*-23 | B4PU4NBNB | B4 | AT | 3UP4S>4I | AUTBS>4I | AUT4S>4I | AUT4S>4I | AUBOS>4I | BUBNS>4I | 3U4OS>4I | | | | NI | NT | 334 | NJ | 3JJ | TJ | | | | 0#-.%8-*%&' | 3AIJONU4B |
| !$(P'C | EJ&? | EJ&? | EO4:/:=N-3/42N*,/8347*:2+/H4<9=938,4? | #C"A#$@@% | D@ | D@ | #A#?EFCB | #A?DEFCB | "A$$EFCB | #A#?EFCB | #A"@EFC> | #AB?EFC> | BA%DEFC> | | | | DBB | D%D | "?? | B% | @> | B@ | | | | GH81-,8*34I. | D%>?CAB? |
| (4II4O | /B( | /!(0^ | /<F"#)-<8#=F8#2B21=&$1=-*#+8#=F8)&6#'-$# | ATPUBITN | A3 | A4 | 3UTIS>4J | IU3BS>4I | AU3OS>4I | IU3BS>4I | OUJOS>4I | AU34S>4I | AUBOS>4I | AUPNS>4B | BUNAS>4B | IUONS>4B | APNB | 3NBJ | 3ABN | 3IIP | IOP | A34 | 3TP | AJO | BIA | 0#-.%8-*%&' | BO4JBUA4P |
| !@(>)" | JJRS"4IEJ&BI | JJRS"4IEJ&BI | J/:=N-3/4:-.3/<89:4,N-3<2*N,/N4" | D$"A$$"B | #C | #C | "A??EFCB | "ABBEFCB | "ACDEFCB | "ABBEFCB | BA#?EFC# | "A"%EFC> | DA$>EFC> | | | | D>> | D@C | "$" | D$ | B# | @" | | | | GH81-,8*34I. | ">%@BA?@ |
| !"BCC% | EJ&D | EJ&D | EO4:/:=N-3/42N*,/8347*:2+/H4<9=938,4D | D??AB%B | D> | D> | $A#@EFCB | %A@$EFCB | @A@%EFCB | @A@%EFCB | "A%"EFCB | "A@@EFCB | "A$?EFCB | | | | #?$ | >C> | >D> | ?D | @@ | "C% | | | | '7/,6+-,8*34I | #>@##A |
| (4I4PJ | !50E! | !50E! | _);"*&$#2D%$1=&$1=-*#+-<8&<-$#+! | APPUB4III | 3P | I | 3UIBS>4J | IUAOS>4I | AU4JS>4I | IUAOS>4I | JUABS>4I | JUTNS>4B | JUJBS>4A | JUOAS>4B | OUNOS>4B | OUPAS>4B | ONT | APA | 3PI | BBP | JO | NI | PN | PA | 34T | [-)D-.%8&.#* | BTIA4U43 |
| !$PCTD | EJ&# | EJ&# | EO4:/:=N-3/42N*,/8347*:2+/H4<9=938,4# | D?#A$B$D | D@ | D@ | DAB$EFCB | "A$@EFCB | "A%?EFCB | "A$@EFCB | "A$%EFC> | DA$DEFC> | BACBEFC> | | | | DDD | DB# | "@? | D> | B# | @@ | | | | '7/,6+-,8*34I | D$$B"A$? |
| G>#>CD | EJ&@ | EJ&@ | EO4:/:=N-3/42N*,/8347*:2+/H4<9=938,4@ | D%"A>>$$D | "? | "? | ?A#%EFC> | %A#@EFC> | BA#%EFC> | %A#@EFC> | "ACDEFC# | "ADDEFC# | ?A?>EFC# | | | | "C% | ""@ | @@ | D? | D? | D" | | | | M/-:81-,8*34 | D#??#AC## |
| !BU&&> | EJ&"C | EJ&"C | EO4:/:=N-3/42N*,/8347*:2+/H4<9=938,4"C | DB"AB%B# | "% | "% | >A#CEFC> | DA$#EFC> | "A@?EFC> | DA$#EFC> | "A>?EFC> | "A@@EFC> | DABBEFC> | | | | "## | ">% | ""? | >C | D$ | D> | | | | M/-:81-,8*34 | D?#>%A@I |
| (40(AJ | [!5CB | [!5CB | [-<.&8;<%'2B | AIOUO4J4B | 33 | 33 | OUJOS>4I | BUJ4S>4I | IUN4S>4I | IUN4S>4I | AU4IS>4I | 3U3IS>4I | IUN4S>4I | IU3NS>4B | | | 3BT | 34J | T3 | JP | AN | 333 | A4 | | | !"#*F<-*%&'- | 3ONBPUJN. |
| (40(AI | [!5CA | [!5CA | [-<.&8;<%'2A | AIOUO4J4B | 33 | 33 | OUJOS>4I | BUJ4S>4I | IUN4S>4I | IUN4S>4I | AU4IS>4I | 3U3IS>4I | IUN4S>4I | IU3NS>4B | | | 3BT | 34J | T3 | JP | AN | 333 | A4 | | | !"#*F<-*%&'- | 3ONBPUJN. |
| (40(AB | [!5C3 | [!5C3 | [-<.&8;<%'23 | AIOUO4J4B | 33 | 33 | OUJOS>4I | BUJ4S>4I | IUN4S>4I | IUN4S>4I | AU4IS>4I | 3U3IS>4I | IUN4S>4I | IU3NS>4B | | | 3BT | 34J | T3 | JP | AN | 333 | A4 | | | !"#*F<-*%&'- | 3ONBPUJN. |
| !BV@J# | EJ&> | EJ&> | EO4:/:=N-3/42N*,/8347*:2+/H4<9=938,4> | D"#ADC%#@ | "# | "# | #AC"EFCB | DA"BEFCB | DA"#EFCB | DA"BEFCB | %A"$EFC> | %AD@EFC> | @A%@EFC> | | | | "D% | "D" | "C> | >$ | >% | >D | | | | M/-:81-,8*34 | DCC@%AB |
| (4J34T | G34!N | G344!N | ()&*#%'+G3442!N | 3O4UJJOA3 | J | J | OUO3S>4I | IUIBS>4I | JUN4S>4I | JUN4S>4I | OUIAS>4I | JUIBS>4I | 3UBIS>4J | AUBNS>4B | 3U3BS>4I | 3U3AS>4I | AIT | A4P | ANI | II | IJ | IT | AJ | AJ | 3A | 0#-.%8-*%&' | 34NBIUJ4O |
| !$Q)@" | EJ&% | EJ&% | EO4:/:=N-3/42N*,/8347*:2+/H4<9=938,4% | ">CA>>$%$ | > | > | @A"DEFC> | ?AC$EFC> | %AB#EFC> | ?AC$EFC> | "A$$EFC> | DA##EFC> | BA%@EFC> | | | | B> | #? | >" | D$ | "B | DB | | | | &-N=-:81*:/, | "DC"?AD$" |

5/./31
2345&6V(X+`+$%6'%a%"-'"#+-$+"-<";<-*#8+DF+2345&6V(X)Q+#X
7(#1*%8#$+`+*&*-<+';.D#)+1#1*%8#$+a&)+*=#+1)&*#%'+8#*#"*#8+-")&$$+-<<+#Z1#)%.#'*$+%'+8-*-$#*+V.&8%a%"-*%&'+8&#$+.-**#)X
79'%:;#+`+*&*-<+';.D#)+&a+;'%:;#+1#1*%8#$+a&)+*=#+1)&*#%'+8#*#"*#8+-")&$$+-<<+#Z1#)%.#'*$+%'+8-*-$#*+b;'%:;#+8#a%'#8+-$+1#1*%8#$+*=-*+-)#+-**)%D;*#8+*&+-+$1#"%a%"+$#:;#'"#+V.&8%a%"-*%&'+8&#$+'&*+.-**#)X
@+`+@#1<%"-*#
!)#-+`+!)#-+;'8#)+*=#+CG3+$1#"*);.+";)M#+*=-*+"-'+D#+-**)%D;*#8+*&+1#1*%8#$+*=-*+-)#+;'%:;#+*&+*=-*+1)&*#%'
7G1#"+`+G1#"*)-<+[&;'*$+-**)%D;*#8+*&+*=-*+1)&*#%'
(LC+`+1&$*2*)-'$<-*%&'-<+.&8%a%"-*%&'$+8#*#"*#8+&'+1#1*%8#$+-**)%D;*#8+*&+*=-*+1)&*#%'

**Table S1 Proteins identified by mass spectrometry**. Filters were applied as described in experimental methods. (See also separate Excel spreadsheet)

#### Table S2 Statistics for data collection, refinement, and validation

| | $Ca_V1.2(\Delta C)/Ca_V\beta_3$ | $Ca_V1.2(\Delta C)/Ca_V\beta_3/Ca_V\alpha_2\delta$-1 | |
|---|---|---|---|
| **Data collection and processing** | | | |
| Magnification | 105,000 | 105,000 | |
| Voltage (kV) | 300 | 300 | |
| Electron dose (e-/Å$^2$) | 46 | 46 | |
| Defocus range (μm) | -0.9~-1.7 | -0.9~-1.7 | |
| Pixel size (Å) | 0.8466 | 0.8466 | |
| Symmetry | C1 | C1 | |
| Initial particle images (no.) | 3,096,005 | 4,522,745 | |
| Final particle images (no.) | 217,117 | 269,950 | 269,802 |
| Map resolution (Å) | 3.4 | 3.6 | 3.3 |
| FSC threshold | 0.143 | 0.143 | 0.143 |
| Map resolution range (Å) | 3.0~8.0 | 3.0~8.0 | 2.0~4.8 |
| **Refinement** | ECAB Map 3 **(PDB:8EOI;EMD-28376)** | | CABAD Map 2 **(PDB:8EOG;EMD-28375**) |
| Initial model used (PDB code) | 6WW7, 7MIY | | 7MIY |
| Model resolution (Å) | 3.4 | | 3.5 |
| FSC threshold | 0.5 | | 0.5 |
| Map sharpening *B* factor (Å$^2$) | -27.6 | | -82.1 |
| Model composition | | | |
| Non-hydrogen atoms | 28,721 | | 19,757 |
| Protein residues | 3,562 | | 2,416 |
| Ligands | 8 | | 24 |
| *B* factors (Å$^2$) | | | |
| Protein | 176.18 | | 129.67 |
| Ligand | 105.71 | | 75.97 |
| R.m.s deviations | | | |
| Bond lengths (Å) | 0.004 | | 0.004 |
| Bond angles (°) | 0.538 | | 0.568 |
| Validation | | | |
| MolProbity score | 2.24 | | 2.5 |
| Clashscore | 8.85 | | 10.78 |
| Poor rotamers (%) | 2.79 | | 3.97 |
| Ramachandran plot | | | |
| Favored (%) | 93.5 | | 91.77 |
| Allowed (%) | 6.47 | | 8.1 |
| Disallowed (%) | 0.03 | | 0.13 |
